# Selective gating of retinal information by arousal

**DOI:** 10.1101/2020.03.12.989913

**Authors:** Liang Liang, Alex Fratzl, Omar El Mansour, Jasmine D.S. Reggiani, Chinfei Chen, Mark L. Andermann

## Abstract

How sensory information is processed by the brain can depend on behavioral state. In the visual thalamus and cortex, arousal/locomotion is associated with changes in the magnitude of responses to visual stimuli. Here, we asked whether such modulation of visual responses might already occur at an earlier stage in this visual pathway. We measured neural activity of retinal axons using wide-field and two-photon calcium imaging in awake mouse thalamus across arousal states associated with different pupil sizes. Surprisingly, visual responses to drifting gratings in retinal axonal boutons were robustly modulated by arousal level, in a manner that varied across stimulus dimensions and across functionally distinct subsets of boutons. At low and intermediate spatial frequencies, the majority of boutons were suppressed by arousal. In contrast, at high spatial frequencies, the proportions of boutons showing enhancement or suppression were more similar, particularly for boutons tuned to regions of visual space ahead of the mouse. Arousal-related modulation also varied with a bouton’s sensitivity to luminance changes and direction of motion, with greater response suppression in boutons tuned to luminance decrements vs. increments, and in boutons preferring motion along directions or axes of optic flow. Together, our results suggest that differential filtering of distinct visual information channels by arousal state occurs at very early stages of visual processing, before the information is transmitted to neurons in visual thalamus. Such early filtering may provide an efficient means of optimizing central visual processing and perception of state-relevant visual stimuli.

## Results

### Bulk visual responses in retinothalamic axons are suppressed by arousal

To test whether arousal states modulate visual responses in retinal ganglion cell (RGC) axons projecting to the dLGN, we first conducted widefield calcium imaging of RGC axons in awake, head-fixed mice (Figure 1A). We presented full-field drifting sinusoidal gratings to the contralateral eye as described previously [1], and used pupil area to assess instantaneous arousal levels (Figure 1B), similar to previous studies [2–6]. GCaMP6f-expressing retinal axons were visualized through a cranial window that was chronically implanted over the thalamus [1](Figure 1C). The most responsive region within the dLGN was selected as the region of interest (ROI) to quantify visual responses across arousal states. Across individual presentations of the identical visual stimulus, the evoked response of the population of RGC axons showed a strong negative correlation with pupil area (e.g. Figure 1D and S1A-C) that was not observed in a control ROI outside the dLGN, in a region that lacked RGC axons (Figure S1D-F). We defined ‘low arousal’ or ‘high arousal’ trials as those with pupil area in the lower or upper 25%, respectively, of the distribution of pupil areas across all trials recorded during a given session (Figure 1D, shaded regions). In the example ROI in Figure 1C-E, the mean amplitude of responses to visual stimuli of an intermediate spatial frequency (0.08 cycles per degree [cpd]) showed an approximately two-fold decrease during high vs. low arousal. To quantify the degree of response modulation, we used a modulation index (MI; [R_HighArousal_ – R_LowArousal_]/R_AllTrials_; see Methods) for which negative values indicate a suppression of the magnitude of visual responses during high arousal trials as compared to low arousal trials. For the above example, this corresponded to a modulation index of −1.14. We observed similar suppression of responses to this visual stimulus in all six mice imaged in this manner (Figure 1G, middle column).

**Figure 1.**
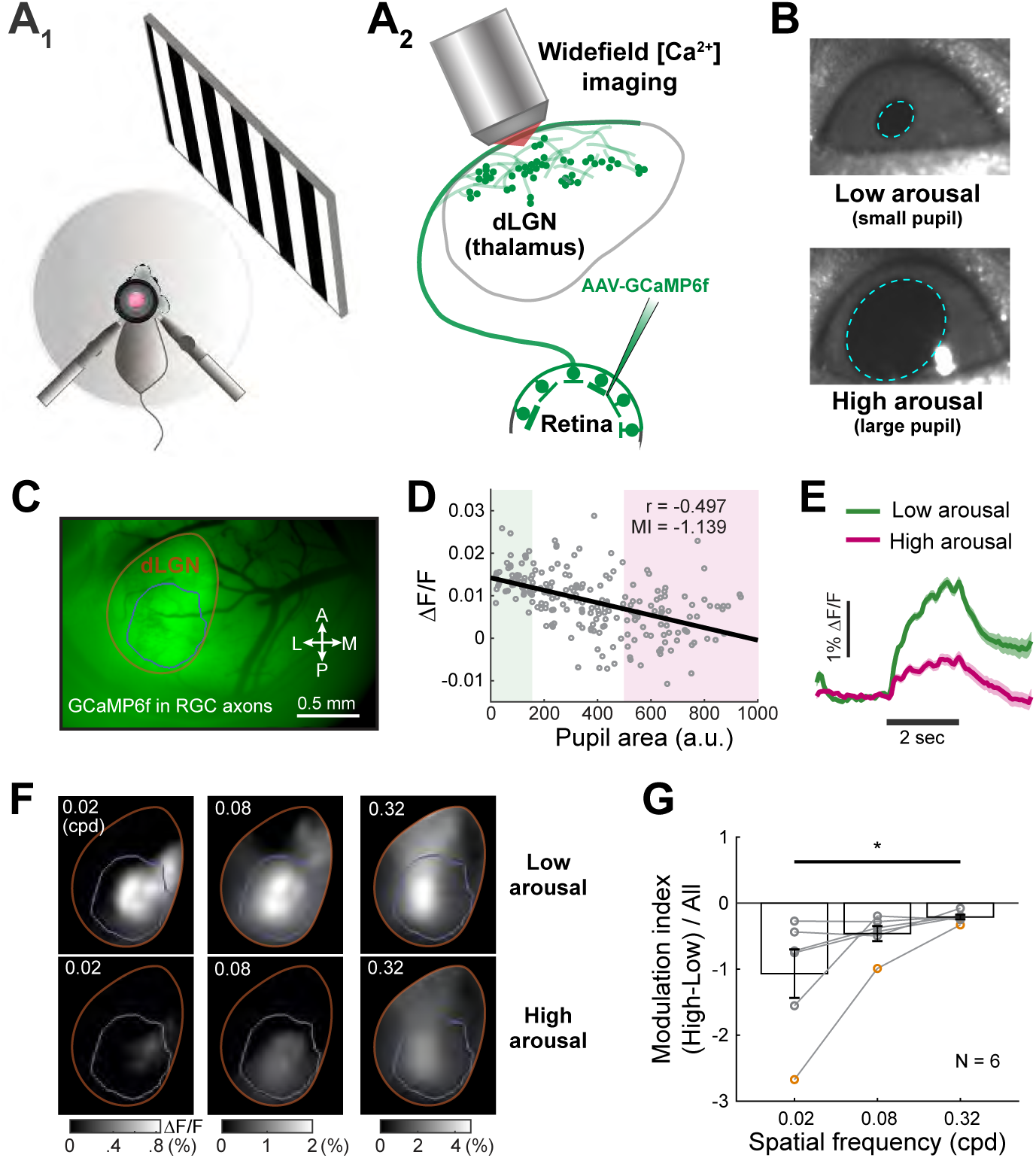
Suppression of bulk visual responses in retinal axons in dLGN during arousal. **A. A_1_**, Setup for imaging visual responses from an awake, head-fixed mouse. **A_2_**, Calcium signals in RGC axons were visualized using widefield epifluorescence imaging. AAV-GCaMP6f was previously injected in the retina to infect retinal ganglion cells. **B.** Arousal levels were estimated by pupil area (outlined in cyan). **C.** Example image of raw fluorescence from retinal axons in the dLGN (outlined in orange) through a chronically implanted cranial window. GCaMP6f was expressed in retinal ganglion cells (RGCs). The region of interest (ROI) used for estimation of bulk activity is outlined in blue. A, P, M, L: anterior, posterior, medial, lateral. **D.** Single-trial visual responses to full-field sinusoidal drifting gratings (direction: nasal to temporal; spatial frequency: 0.08 cycles per degree [cpd]) were negatively correlated with pupil area in the example recording in C (r = −0.497, p < 10-14). The region encompassing the 25% trials with the smallest pupil area (Low arousal trials) was labeled in pale green. The region encompassing the 25% trials with the largest pupil area (High arousal trials) was labeled in pale magenta. ΔF/F: fractional change in fluorescence. a.u.: arbitrary units. **E.** Average time courses of visual responses to drifting gratings of 0.08 cpd during low (green) and high (magenta) arousal states. Mean +/− SEM. **F.** Images of average responses during low and high arousal states across spatial frequencies suggest that visual responses are suppressed by arousal. This suppression was stronger at lower spatial frequencies. Data in C-F are from the same mouse. **G.** Modulation index (MI) across mice (N=6) at 3 different spatial frequencies. Index values were negative in all cases, indicating that arousal consistently suppressed bulk visual responses across retinal axons. Suppression was strongest at lower spatial frequencies (MIs were significantly different between 0.02 cpd and 0.32 cpd, p < 0.05, Friedman test with post-hoc Dunn’s correction). Data from the example mouse in C-F are labeled in orange. Mean ± SEM. N=6 mice.

We next examined how this arousal-dependent response suppression varied across grating stimuli spanning four octaves in spatial frequency (SF; 0.02, 0.08, 0.32 cpd). In the example ROI in Figure 1C-F, arousal led to suppression of responses at all spatial frequencies. This arousal-dependent suppression was observed for RGC axon responses in the dLGN (but not in control ROIs from the same experiments) in 6/6 mice (Figure 1G, S1E and S1D, F). Moreover, suppression was significantly greater at low vs. high spatial frequencies (Figure 1F-G; 0.02 cpd vs. 0.32 cpd), suggesting greater net suppression of larger objects.

Several controls confirmed that arousal-dependent modulation of visual responses in retinothalamic axonal boutons was correlated with changes in pupil area over the timescale of tens of seconds, rather than over longer timescales. Specifically, one potential concern was that while pupil area exhibited large fluctuations across tens of seconds (Figure S1A_1_), it might also exhibit slower changes across tens of minutes. If so, response differences between large and small pupil trials could partially reflect slow and systematic changes in responses across a session. However, inspection of pupil area and response magnitude from the example session in Figure 1C-F demonstrated changes in response magnitude that were inversely proportional with changes in pupil area even in consecutive trials (Figure S1A_1_). Further, the inverse correlation between response magnitude and pupil area was consistent when considering the first and second half of the session separately (Figure S1B). We additionally confirmed that the observed arousal-dependent suppression was not related to any cue-evoked changes in pupil diameter, which were either negligibly small or absent (Figure S1C).

### Arousal-dependent response suppression in individual retinal boutons in the dLGN

The above results involving bulk recording of axons led us to ask: does arousal cause suppression in all retinal axonal boutons, or only in specific subsets of boutons? We therefore recorded simultaneous calcium signals from hundreds of retinal boutons in the ‘shell’ of the dLGN of awake mice using two-photon calcium imaging (Figure 2A-C; ∼20-90 μm below the surface of the dLGN or 80-150 um below the surface of the optic tract)[1], while monitoring pupil size and running speed to characterize states of arousal and locomotion (Figure S2A-B). As shown for an example direction-selective RGC bouton, the magnitude of responses to a given stimulus varied across individual trials (Figure 2D). Response amplitudes were negatively correlated with pupil area (Figure 2E), and single-trial responses during low arousal displayed higher response magnitudes than during high arousal (Figure 2F). A similar degree of arousal-dependent suppression was evident not only for stimuli presented at the bouton’s preferred direction of motion, but also across all other directions sampled (Figure 2G, S2C). In this way, this multiplicative gain change with arousal did not strongly affect direction preference or direction selectivity for this example bouton (Figure 2G) or for most other direction-selective boutons (Figure S2D-E; cf. [7]). Similar arousal-dependent suppression was observed in the majority of direction-selective boutons (4324 boutons from 22 FOVs in 5 mice; Figure S2F), both at the level of correlation coefficients and of modulation index values (Figure 2H-I). These arousal-dependent changes in the magnitude of the evoked response (⊗F/F_0_) could not be accounted for by the very small and uncorrelated changes in pre-stimulus baseline fluorescence, F_0_ (Figure S2G-I).

**Figure 2.**
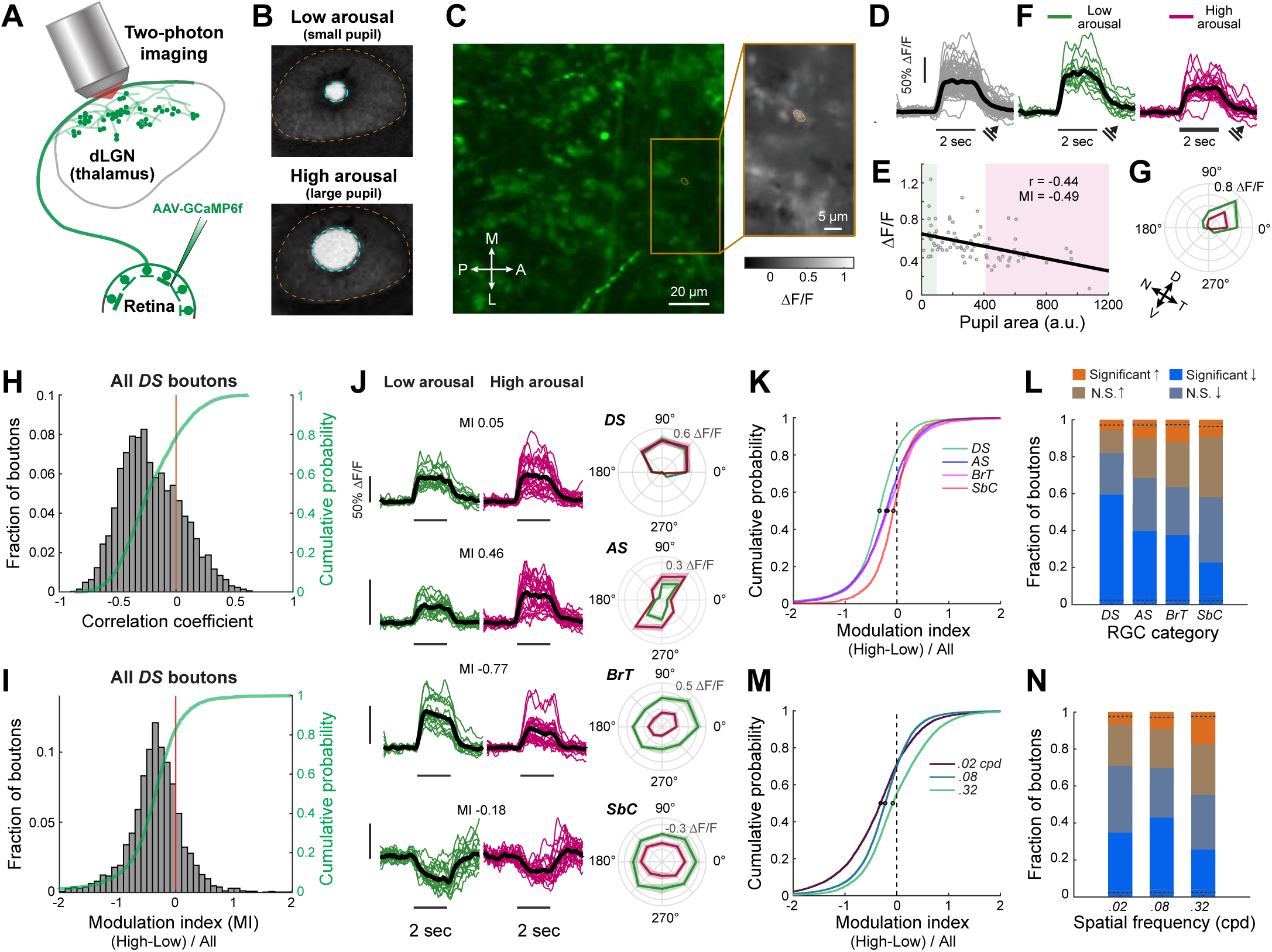
Arousal suppresses visual responses in individual retinal boutons in the dLGN. **A.** Setup for two-photon calcium imaging of visual responses of RGC axons expressing GCaMP6f in awake mice. **B.** Arousal levels were estimated by pupil area. The pupil (outlined in cyan) of the eye (outlined in orange) was illuminated by transmission of infrared laser light entering the cranial window and passing through the brain. **C.** Left: example raw fluorescence image of retinal axons and axonal boutons in the dLGN, 115 m below the window. M, medial. L, lateral. A, anterior. P, posterior. Right: average visually evoked calcium response in RGC boutons from a subregion outlined in the image at left. ΔF/F: fractional change in fluorescence. **D.** Reliable single-trial visual responses of an individual RGC bouton (outlined in orange in C, right) to repeated presentations of the same grating stimulus, drifting along the bouton’s preferred direction (45°, see arrow; spatial frequency: 0.08 cpd). **E.** Single-trial visual response magnitudes in D were negatively correlated with pupil area (r = −0.44, p < 10-4, 95%confidence interval for r from shuffled data: [-0.209, 0.225]). Low and High arousal states were shaded in pale green and magenta, respectively. **F.** Single-trial response time courses for this bouton during states of low arousal (green) and high arousal (magenta). **G.** Average direction tuning curves of the same RGC bouton as in D, at low arousal (green) and high arousal (magenta). In Figures 2-3, 0° indicates rightward (roughly nasal-to-temporal) motion on the LCD monitor, and 90°indicates upward motion. Arrows: estimated nasal-temporal and dorsal-ventral axes, if the head were in typical ambulatory posture. N, nasal movement. T, temporal. D, dorsal. V, ventral. Mean ± SEM. **H.** Visual responses evoked by drifting gratings of 0.08 cpd were negatively correlated with arousal in the majority of the direction-selective (DS) boutons (78.3%, 3386/4324 boutons from 22 FOV from 5 mice). Median correlation, r = −0.246. Red line, r = 0. Green: cumulative distribution. **I.** The majority of DS boutons (82.1%, 2971/3619 boutons from 22 FOV from 5 mice), had negative modulation indices (MIs; see STAR Methods).Median correlation: r = −0.336. **J.** Single-trial responses and direction tuning curves during low (green) and high (magenta) arousal, from example direction-selective (DS), axis-selective (AS), broadly tuned (BrT) and Suppressed-by-contrast (SbC) boutons. Note that the example BrT and SbC boutons were suppressed by arousal, while the example AS bouton was enhanced by arousal. **K.** Cumulative distributions of modulation indices for different bouton categories (N = 3619 DS, 3609 AS, 3692 BrT, and 1521 SbC boutons). Median values are circled in black. For DS, AS, BrT and SbC boutons, median values were −0.336, −0.208, −0.182, −0.069, respectively. 82.1%, 68.6%, 63.6%, 58.2%of the boutons had negative MI values, respectively. MI values in K-L were derived from responses to gratings of 0.08 cpd. The cumulative distributions between bouton categories were significantly different from each other (p’s < 10-4, Kolmogorov-Smirnov tests with Bonferroni correction). **L.** Fraction of boutons per category that were significantly enhanced, not significantly (N.S.) enhanced, not significantly suppressed, or significantly suppressed. Black dashed lines indicate the fraction of boutons that were significantly enhanced or significantly suppressed in shuffled data (median across 1000 shuffles). Gray lines indicate associated 95% confidence intervals. **M.** Cumulative distributions of modulation indices for different spatial frequencies, pooled across bouton categories (N = 8074, 12441, and 11311 boutons with significant responses at 0.02, 0.08 and 0.32 cpd, respectively). Median values were −0.314, −0.224 and −0.077, and 71.0%, 69.8%, 55.3% of the boutons had negative MI values, respectively. Distributions for different spatial frequencies were all significantly different from each other (all p’s < 10-63, Kolmogorov-Smirnov tests with Bonferroni correction). **N.** Fraction of boutons driven at a given spatial frequency that were significantly enhanced, not significantly (N.S.) enhanced, not significantly suppressed, or significantly suppressed. Black dashed lines indicate the fraction of boutons that were significantly enhanced or significantly suppressed in shuffled data (median across 1000 shuffles). Gray lines indicate associated 95% confidence intervals.

We have previously demonstrated that the shell of mouse dLGN receives inputs from functionally diverse types of retinal ganglion cells, which we characterized into four broad functional categories: direction-selective (DS; selective to one direction of motion, see above), axis-selective (AS; responsive to opposite directions of motion along the same axis, also known as ‘‘orientation-selective’’), broadly-tuned (BrT; broadly responsive across all directions), and suppressed-by-contrast (SbC) [1]. Boutons in each category could be significantly enhanced, suppressed, or unaltered by arousal (Figure 2J-L). The majority of boutons in each category had responses that were suppressed by arousal (Figure 2K). Suppression by arousal was statistically significant in a large proportion of individual boutons, while a substantially smaller proportion of boutons were significantly enhanced by arousal (Figure 2L). Notably, while locomotion is known to correlate with enlarged pupil size [3,5,8,9], arousal level could also fluctuate across stationary trials (Figure S2A). We confirmed that the above findings persisted even when the subset of trials involving locomotion were excluded (Figure S2J-L).

Consistent with bulk recordings from retinal axons (Figure 1), the population of individual RGC boutons showed greater arousal-dependent suppression of responses to low vs. high spatial frequency stimuli (Figure 2M). Accordingly, a larger proportion of boutons showed significant arousal-related response suppression for low/intermediate vs. high spatial frequency stimuli (Figure 2N). Similarly, a smaller proportion showed significant response enhancement for low/intermediate vs. high spatial frequency stimuli (Figure 2N).

### Greater arousal-related response suppression in ‘Off’ vs. ‘On’ retinal boutons

The above findings show that arousal-dependent modulation of visual responses is a common feature of RGC boutons in the dLGN, and that this modulation depends on stimulus spatial frequency. However, even at a given spatial frequency, substantial heterogeneity exists across boutons in each broad RGC category (Figure 2L). Thus, we wondered whether a bouton’s arousal-dependent response modulation also varied with other tuning properties. First, we considered each bouton’s responses to full-field luminance changes, and classified boutons as either ‘On’ type RGCs (selective to luminance increases), ‘Off’ type RGCs (selective to luminance decreases), or ‘On-Off’ type RGCs (responding to both) (Figure 3A-B). For each bouton, estimated the arousal modulation index using the responses to the same full-field drifting grating stimuli as above (Figure 3A). Similar numbers of On boutons were enhanced or suppressed by arousal, while ∼80% of Off boutons were suppressed by arousal, and modulation index values differed significantly between On and Off boutons (Figure 3C-D). We observed similar differences in arousal modulation between On and Off boutons when we separated these boutons into each of the four broad RGC categories (Figure S3A-B), and when we considered responses to stimulation at other spatial frequencies (Figure S3C-F).

**Figure 3.**
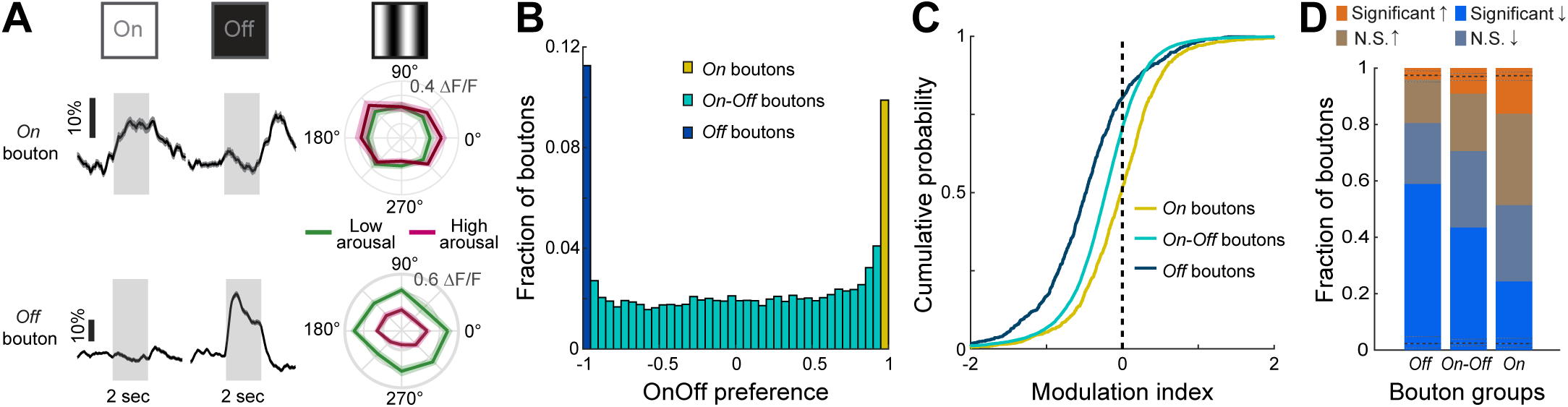
Greater arousal-related response suppression in Off vs. On retinal boutons. **A.** Average response time courses during full-field luminance increments (left) and decrements (middle), and direction tuning curves of average responses to full-field drifting gratings (right), from an example On bouton and an example Off bouton. Time courses and tuning curves: mean ± SEM. **B.** Fraction of boutons defined as ‘On’ (OnOff preference index > 0.95, n =1364), ‘Off’ (OnOff preference index < −0.95, n = 1553) or ‘On-Off’ (OnOff preference index within [-0.95, 0.95], n = 10877). Data from 22 FOV from 5 mice. **C.** Cumulative distributions of modulation indices for On, On-Off, and Off boutons. Distributions were all significantly different from each other (all p’s < 10-34, Kolmogorov-Smirnov tests with Bonferroni correction). Median index value for On, On-Off and Off boutons: −0.016, −0.222, −0.479, respectively. 51.5%, 70.6% and 80.5% of On, On-Off, and Off boutons had negative index values, respectively. Indices were calculated from responses to drifting gratings at 0.08 cpd. **D.** Fraction of boutons of each type that were significantly enhanced, not significantly (N.S.) enhanced, not significantly suppressed, or significantly suppressed. Black dashed lines indicate the fraction of boutons that were significantly enhanced or significantly suppressed in shuffled data (median across 1000 shuffles). Gray lines indicate associated 95% confidence intervals. Indices were calculated from responses to drifting gratings at 0.08 cpd.

### State-dependent suppression depends on direction and axis preference

The significance of visual motion along different directions or axes might change depending on behavioral state. In the case of an unaroused and stationary mouse, detection of any motion may be essential to respond to approaching predators or other threats. In contrast, for an aroused mouse that is moving forward, visual motion signals will include self-generated optic flow signals. We thus considered arousal modulation of responses to moving gratings in two non-overlapping categories of retinal boutons that were either tuned to a specific motion direction (direction selective, DS) or to a specific motion axis (axis selective, AS). We previously showed that the subregion of dorsal dLGN that we consistently targeted across mice for imaging (corresponding to RGC boutons with dorsal and somewhat lateral visual field preferences) contains DS boutons that prefer one of four directions of motion. The majority of these DS boutons prefer posterior or upward motion in visual space (Liang et al., 2018; Figure 4A). While all four groups showed arousal-dependent suppression of responses to drifting gratings, suppression was larger and more common for boutons preferring posterior or upward motion. This directional bias in arousal modulation may serve to attenuate self-generated visual signals related to optic flow, which is anticipated during current or upcoming locomotion, while preserving encoding of other motion in this region of visual space that might indicate motion of animate objects. Similar findings were observed across a range of spatial frequencies (0.02-0.32 cpd; Figure S4A-B). We also analyzed axis-selective (AS) boutons tuned to motion along a specific axis. AS boutons preferring motion roughly along the horizon were more commonly suppressed than AS boutons preferring vertical motion (Figure 4B and S4C-D).

**Figure 4.**
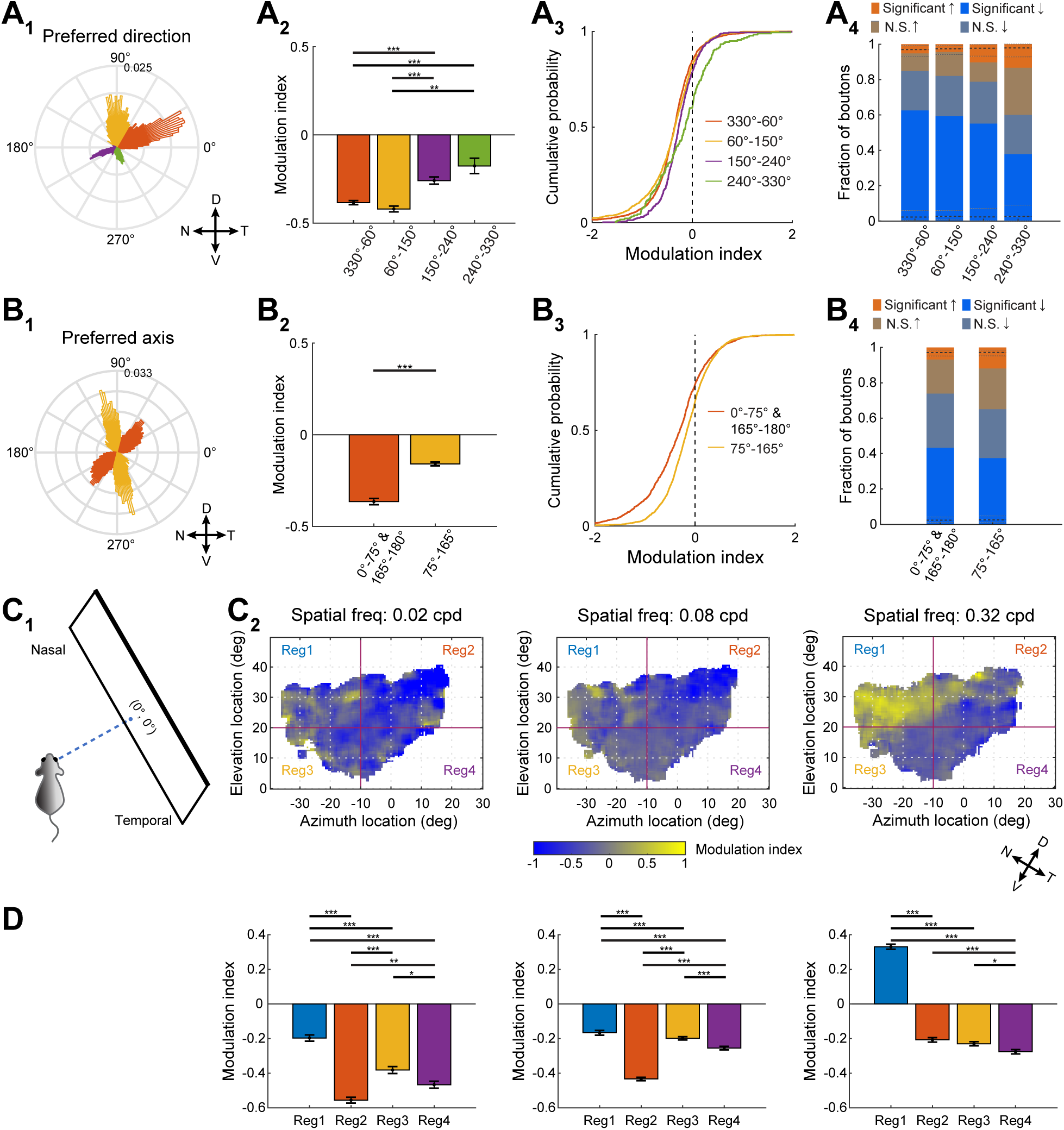
Differential arousal modulation of boutons with distinct motion preferences or retinotopic preferences. **A. A_1_**, distribution of the preferred motion direction across all direction-selective (DS) boutons recorded in dorsal dLGN. DS boutons can be roughly divided into 4 groups based on the four peaks in the overall distribution of direction preferences. We therefore defined these groups manually, and labeled each group with a distinct color. Ns for groups of boutons with direction preferences between 330°-60°, 60°-150°, 150°-240°, and 240°-330° were 1924, 1143, 372, and 180, respectively. Data from 22 FOV from 5 mice. **A_2_,** average arousal modulation index of DS boutons in each group. Mean ± SEM. Median value for each group: −0.356, −0.350, −0.272, −0.137. Distributions of indices were significantly different across groups (p < 10-8, Kruskal-Wallis test with Dunn correction; ***p < 0.001). **A_3_**, cumulative distributions of modulation indices for each group. Indices were negative for 84.8%, 82.1%, 78.8%, and 60.0% of boutons in each group. Distributions were all significantly different from each other (all p’s < 10-3, Kolmogorov-Smirnov tests with Bonferroni correction), except for between 330°-60° and 60°-150° (p = 0.10) or between 150°-240° and 240°-330° (p = 0.0015). **A_4_,** Fraction of boutons per group that were significantly enhanced, not significantly (N.S.) enhanced, not significantly suppressed, or significantly suppressed. Black dashed lines indicate the fraction of boutons that were significantly enhanced or significantly suppressed in shuffled data (median across 1000 shuffles). Gray lines indicate associated 95% confidence intervals. Indices were calculated from responses to drifting gratings at 0.08 cpd. D, V, N, T: dorsal, ventral, nasal, temporal. Note that polar axes have been rotated by 31° such that 0° indicates nasal-to-temporal motion during typical ambulatory posture [29]. **B. B_1_,** distribution of preferred axis of motion across all axis-selective (AS) boutons recorded in the dorsal dLGN. AS boutons can be roughly divided into 2 groups, each labeled by a distinct color. Ns for groups of boutons with preferences ranging from 0°-75° & 165°-180° and 75°-165° were 1402 and 2207, respectively. Note that the distribution of axis preferences from 180°-360° is duplicated from that used for 0°-180° to illustrate that the orange group spans −15°-75° or, equivalently, 0°-75° & 165°-180°. Data from 22 FOV from 5 mice. B_2_, average arousal modulation index of AS boutons in each group. Mean ± SEM. Median value for each group: −0.305, −0.160. Distributions of indices were significantly different across groups (***p < 10-22, Wilcoxon rank sum test). **B_3_**, cumulative distributions of modulation indices for each group. Indices were negative for 74.0% and 65.1% of boutons in each group. Distributions were significantly different (p < 10-22, Kolmogorov-Smirnov test). **B_4_**, Fraction of boutons per group that were significantly enhanced, not significantly (N.S.) enhanced, not significantly suppressed, or significantly suppressed. Black dashed lines indicate the fraction of boutons that were significantly enhanced or significantly suppressed in shuffled data (median across 1000 shuffles). Indices were calculated from responses to drifting gratings at 0.08 cpd. **C. C_1_,** schematics of the relative location of the monitor and the eye contralateral to the imaged boutons. The intersection point between the monitor and an imaginary dashed line that is perpendicular to both the mouse face and the monitor is designated as 0° in azimuth and 0° in elevation. **C_2_,** modulation indices of individual boutons mapped to their preferred retinotopic locations in the visual space. The maps include all DS, AS, BrT and SbC boutons. Each map involves stimuli presented at a different spatial frequency. Maps were smoothed using a 3° window. Portions of the maps of visual space for which no recorded boutons with corresponding retinotopic preferences exist are shown in white. We further divided each map into four regions (see red dividing lines) for comparing MI distributions across regions (see panel D and Figure S5B). Note that the above estimates of elevation and azimuth are in LCD monitor coordinates. To estimate how this relates to natural elevation and azimuth axes during a typical ambulatory posture [29], we plotted arrows at bottom right indicating a rough estimate of nasal-temporal and dorsal-ventral axes. **D.** Average arousal modulation index of the four regions at each spatial frequency. Mean ± SEM. Median value for the four regions at 0.02 cpd: −0.187, −0.532, −0.352, −0.382; at 0.08 cpd: −0.101, −0.391, −0.207, −0.255; at 0.32 cpd: 0.364, −0.237, −0.244, −0.290. Number of boutons for the four regions at 0.02 cpd: 1235, 1872, 1058, 1038; at 0.08 cpd: 1908, 3005, 1730, 1673; at 0.32 cpd: 1654, 2325, 1566, 1433. Distributions of indices were significantly different across regions (p < 10-50, p < 10-102, p < 10-242 for maps at spatial frequencies of 0.02 cpd, 0.08 cpd, and 0.32 cpd, respectively; Kruskal-Wallis test with Dunn correction; ***p < 0.001).

### Arousal-related response suppression depends on retinotopic preference

Does the level and sign of arousal modulation of visual responses vary with a bouton’s retinotopic preference? For the majority of recordings, we also estimated each bouton’s retinotopic preference by presenting elongated bars containing spatiotemporal noise (Figure 4C_1_, Figure S5A, [1]). We then mapped each bouton’s modulation index onto its retinotopic preference in azimuth and elevation, and created a spatial map of arousal modulation for each stimulus spatial frequency (Figure 4C_2_, Figure S5B) and for each functional category of boutons (Figure S5C-F). At the lowest spatial frequency used (0.02 cpd), boutons encoding most regions of visual space showed grating responses that were suppressed by arousal. However, this map became more spatially heterogeneous at the highest spatial frequency used (0.32 cpd), when considering all boutons together (Figure 4C_2_) or when considering each bouton category separately (Figure S5C-F). In this case, arousal actually enhanced responses to high spatial frequency gratings for boutons encoding certain regions of visual space. Specifically, boutons sensitive to stimuli in dorsal and nasal regions of visual space showed, on average, enhanced responses to high spatial frequency stimuli (see Region 1 of Figure 4C_2_, right; Figure 4D). Thus, during arousal, high spatial frequency information from the nasal and dorsal part of the visual field may be enhanced while information from the ventral and lateral parts of the visual field is suppressed.

## Discussion

We demonstrated that visual responses are already modulated by arousal state at a very early stage in the image-forming pathway, before information is transmitted to the visual thalamus. Surprisingly, visually evoked activity in the majority of retinal boutons was suppressed during arousal. This suppression was the prevailing effect in both widefield and two-photon calcium imaging data. Moreover, the extent of suppression depended on the spatial frequency of the visual stimulus, with stronger suppression of responses to lower spatial frequencies. Although the majority of boutons were suppressed by arousal, the responses of some boutons were enhanced, particularly at higher spatial frequencies [10]. Such heterogeneity was evident across all four broadly defined functional categories of RGCs – Direction-tuned, Axis-tuned, Broadly-tuned, and Suppressed. However, when we classified boutons according to preferences for luminance changes, we found that ‘On’ boutons showed equal likelihood of arousal-related enhancement or suppression, whereas ‘Off’ boutons were mostly suppressed by arousal, particularly for categories lacking direction tuning (Broadly-tuned and Suppressed). Arousal modulation also depended on the bouton’s preferred motion direction and motion axis, with greater suppression of responses in boutons tuned to motion along the upward or temporal directions (i.e. directions of optic flow, for neurons coding the region of visual space that we examined). In addition, initial results hint at a topographic map of arousal modulation, with greater suppression in ventral and temporal regions of visual space. Taken together, these findings indicate *differential* modulation of distinct visual information channels prior to the stage of processing by thalamic relay neurons.

Brain state-dependent visual processing has been studied in rodent visual cortical neurons [5,7,10–15] and in dLGN relay neurons [8,9,16]. Recent studies in both primary visual cortex and dLGN have demonstrated significant heterogeneity in modulation by locomotion [8–10,12]. Our findings demonstrating arousal-related response suppression at the level of retinal axon terminals may reflect an efficient and selective means of filtering out irrelevant information before it gets amplified by thalamocortical circuits. This could serve several purposes. First, as described above, the suppression of visual signals related to optic flow may allow more efficient coding of unexpected signals such as those from moving objects in the environment. Second, an increase in arousal may warrant a shift in visual processing from detection of possible threats to high-resolution discrimination of bright objects in front of the animal.

Arousal-related suppression of visual responses observed at the level of RGC axons could be due to several mechanisms, including direct neuromodulation of RGC axons and/or differences in optical filtering of incident stimuli due to variations in pupil diameter. While a dissection of the possible mechanisms involved is beyond the scope of this study, it is worthwhile considering some of these potential causes of modulation. First, a number of neuromodulators are known to act on RGC axon terminals to inhibit presynaptic calcium influx and neurotransmitter release [17–21]. GABA_B_ receptors could mediate direct inhibition at RGC axon terminals from thalamic reticular neurons and / or local inhibitory neurons, both of which receive top-down input from cortex and elsewhere [22, 23]. Interestingly, while serotonin has been shown to inhibit retinothalamic synaptic transmission by activation of 5HT-1B receptors, it can enhance excitability of postsynaptic thalamocortical neurons [24, 25]. This opposing modulation may serve to suppress less relevant signals as early as at the RGC bouton, thus allowing pertinent information to be efficiently transmitted to the dLGN and amplified in downstream circuits.

Finally, it is possible that the increase in pupil diameter drives suppression due to a decreased depth of field [26], or to an increase in overall illumination of the retina. However, these changes are unlikely to explain the majority of our findings. The decreased depth of field during pupil dilation would lead to blurring of stimuli that are presented outside the focal plane, particularly stimuli with higher spatial-frequency content. In contrast, we find the largest arousal-dependent suppression of responses for stimuli with lower spatial frequencies. Similarly, it is not clear whether such optical filtering of the stimulus could explain the dependence of suppression on specific motion directions. Nevertheless, a recent study that observed state-dependent modulation of retinal axonal boutons in superior colliculus also reported modulation of responses using electrophysiology recordings from the optic tract, suggesting that some of the effects of behavioral state might indeed originate in the retina [27], possibly via centrifugal (i.e. retinopetal) projections [28] and/or state-dependent hormonal signals.

Regardless of the potential mechanisms involved, our findings demonstrate that arousal can influence the nature and magnitude of visual information transmission from a very early stage in the image-forming pathway. Behavior states play important roles in shaping neural dynamics, sensory perception and behavior performance. This has led to a surge of studies evaluating state-dependent modulation of visual responses in thalamic and cortical neurons. Our findings highlight the existence of flexible, feature-specific gating of visual responses at even earlier stages of this pathway, providing a powerful, efficient and state-dependent means for exclusion of irrelevant information by the early visual system.

## Acknowledgements

We thank members of the Andermann and Chen labs for helpful discussion. We thank Drs. Jayaraman, Kerr, Kim, Looger, and Svoboda and the GENIE Project, Janelia Farm Research Campus, HHMI, for GCaMP6. We thank the Boston Children’s Hospital viral core for virus packaging. Support was provided by a Simons Collaboration on the Global Brain Postdoctoral Fellowship (to L.L.), a Bertarelli Foundation Fellowship (to A.F.), NIH (R01EY013613 and U54 HD090255 to C.C.), the Harvard/MIT Joint Research Grants Program in Basic Neuroscience and NIH R21 EY030243 (to C.C. and M.L.A.), an NIH DP2DK105570, R01 DK109930, DP1 AT010971, and grants from the Smith Family Foundation, the Pew Scholars Program in the Biomedical Sciences, Klarman Family Foundation (to M.L.A.), the IDDRC Cellular Imaging Core, and the BCH Viral Core (supported by NIH P30 EY012196).

## Author contributions

L.L., A.F., C.C., and M.L.A. conceived and designed the study. L.L. and J.D.S.R. performed surgeries and imaging experiments. L.L., A.F., and O.E.M. analyzed the data. L.L., C.C., and M.L.A. wrote the manuscript with input from all authors.

---

**Figure S1.**
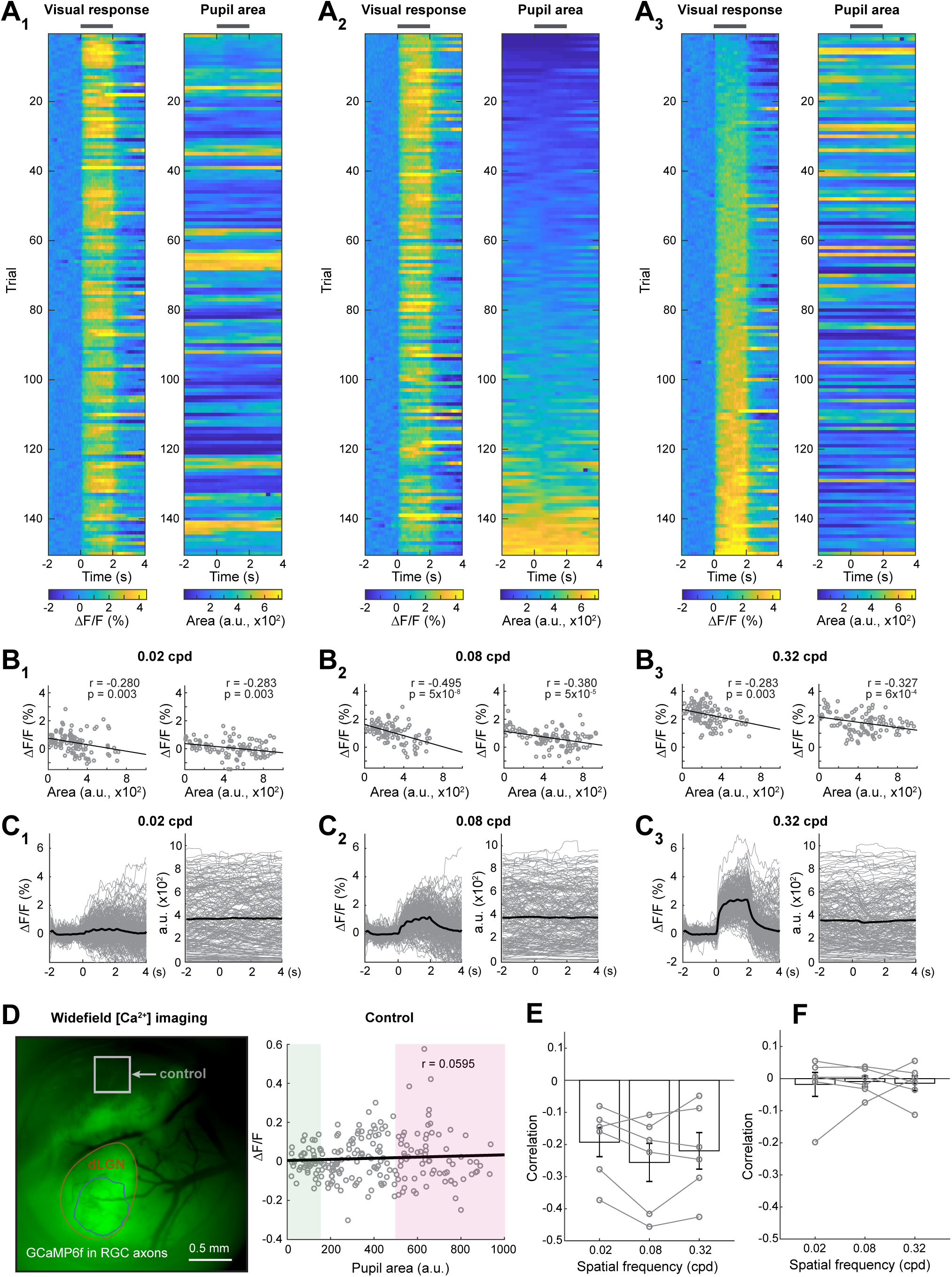
Evoked visual responses are negatively correlated with pupil area, Related to Figure 1. **A.** Example time courses of the evoked visual responses (0.32 cpd gratings) and the corresponding pupil areas, measured from the same mouse as in Figure 1C-F. Dark gray bars, visual stimulus presentation. **A_1_**, time courses from the first 150 trials of the session. **A_2_**, same time courses, re-sorted by average pupil area per trial. **A_3_**, same time courses, re-sorted by the average visual response per trial. ͅF /F, fractional change in fluorescence. **B.** Single-trial mean visually evoked responses (y-axis) vs. pupil area (x-axis) in the first 108 trials (left) or in the second 108 trials (right) of the session in A, for drifting gratings of 0.02 cpd (**B_1_**), 0.08 cpd (**B_2_**), or 0.32 cpd (**B_3_**). Gray discs, single-trial data. Black lines, linear regression fits. r, correlation coefficient. p, p value. **C.** Time courses of calcium responses (left) and pupil areas (right) from all 216 trials, for drifting gratings with spatial frequencies of 0.02 cpd (**C_1_**), 0.08 cpd (**C_2_**), and 0.32 cpd (**C_3_**). Gray, time courses from single trials. Black, mean time courses. Note that visual cues do not drive substantial changes in pupil area within a trial. **D.** Left: location of ROI inside dLGN (outlined in blue) and control ROI outside dLGN (gray square), used for estimating visual responses in an example field of view (FOV). Right: for the control ROI, single-trial visual responses to full-screen drifting gratings (direction: nasal to temporal; spatial frequency: 0.08 cpd) were not correlated with pupil area across trials (r = −0.059; p = 0.384). Low and high arousal states were shaded in pale green and pale magenta, respectively. **E.** Single-trial visual response magnitudes for retinal axons in the ROI located within dLGN were negatively correlated with pupil area for all three spatial frequencies. Mean ± SEM. N=6 mice. **F.** Single-trial visual response magnitudes for the control ROI outside of the dLGN (see panel D) were not correlated with pupil area across trials. Mean ± SEM. N=6 mice. Correlation coefficients from each spatial frequency were not significantly different from each other (Friedman test).

**Figure S2.**
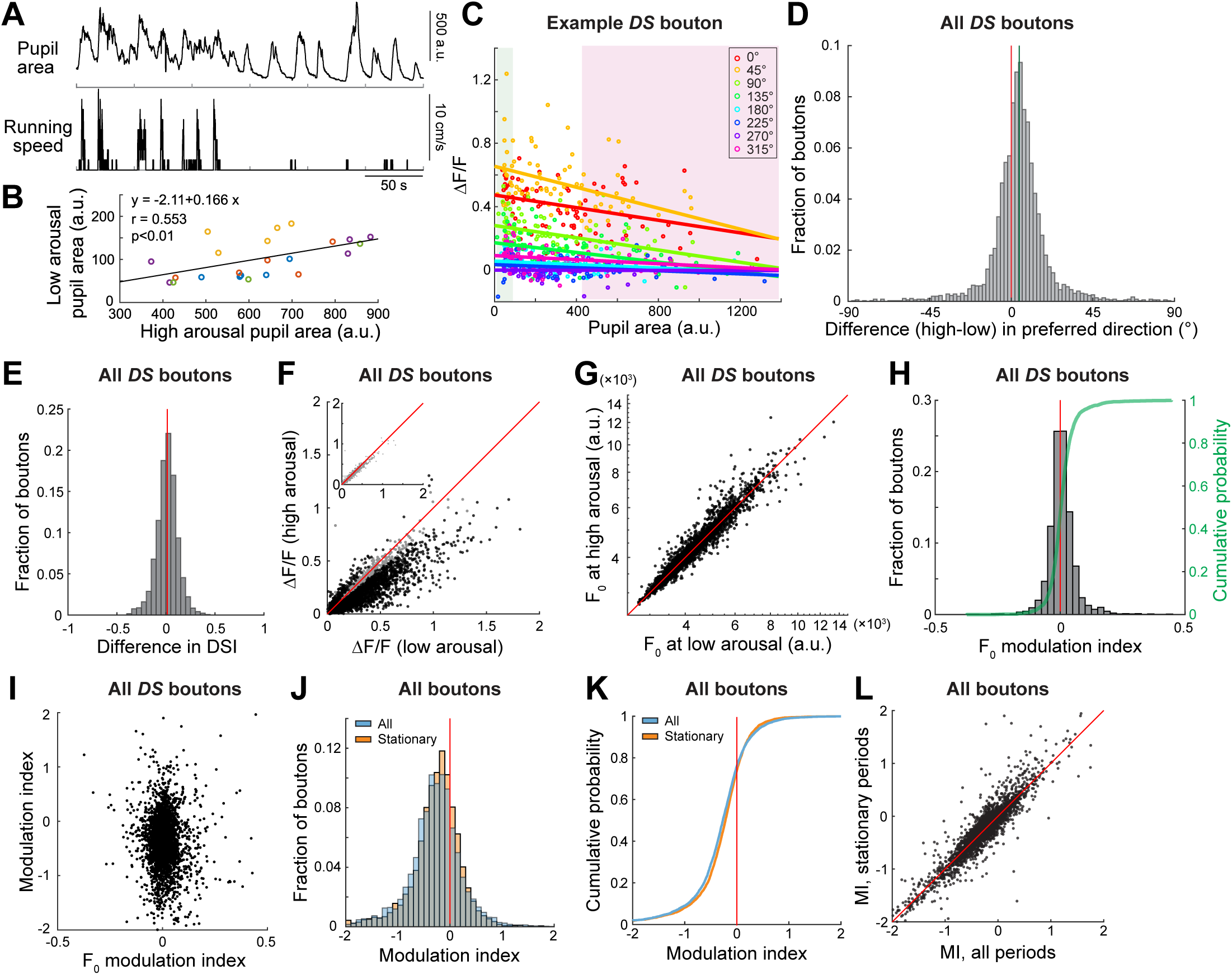
Direction tuning and baseline fluorescence were similar across arousal states; visual responses were similarly modulated by arousal in the absence of locomotion, Related to Figure 2. A. Example epoch with traces of pupil area and running speed. Pupil area fluctuated on a time scale of tens of seconds. During locomotion, pupil area was usually large and covaried with running speed. However, mice could alternate between high and low arousal (i.e. pupil area) even during stationary periods. B. Median pupil area at low arousal vs. high arousal for each experiment/FOV. FOVs from the same mouse share the same color. C. Single-trial visual responses (LiF/F) of the example DS bouton from Figure 2C-G were negatively correlated with pupil area, regardless of the direction of the drifting grating. Trials with the same drifting grating direction share the same color. D. Preferred direction remained largely the same across arousal states. The median value for the difference in preferred direction between high arousal and low arousal states was 4.43° (green line). The differences of the preferred direction were between [-22.5° 22.5°] in 86.6% of the DS boutons and between [-45° 45°] in 94.5% of all the DS boutons. Note that preferred direction was estimated from a fit to a tuning curve generated from stimuli with directions spaced 45° apart. N = 3478 boutons from 22 FOV of 5 mice. Red line: no change in preferred direction. E. Distribution of difference in direction selectivity index (DSI) of DS boutons across states. DSI indices were largely unaffected by arousal state. Median difference in DSI across states: 0.013. N = 3065 boutons. Red line: no change in DSI index. **F.** Scatter plot of mean visually evoked responses at high arousal vs. low arousal across DS boutons. Gray dots in the main plot (also plotted in the inset plot for clarity): boutons that did not show significant differences across arousal states. Black dots: boutons with significantly different response magnitudes between the two states. Responses during low arousal were higher in the majority of the boutons. The evoked responses to drifting gratings at 0.08 cpd were measured at the boutons’ preferred directions. Ntotal = 3619, Nsig = 2364; NN.S. = 1255 boutons from 22 FOV and 5 mice. Red line: line of unity. **G.** Scatter plot of average baseline fluorescence, F0, during high arousal vs. low arousal states, across all DS boutons (N = 3619). P < 10-8, Wilcoxon signed rank test. **H.** The distribution of modulation indices of F0 levels across DS boutons was nearly symmetrically distributed around 0 (red line; N = 3619 boutons from 22 FOV from 5 mice; see STAR Methods). Median: 0.0029. Green: cumulative distribution. 46.6% of DS boutons had negative indices. **I.** The arousal modulation index for evoked responses (ΔF/F, used in Figure 2) was not correlated with the modulation index of F0 across individual boutons. Correlation coefficient: r = −0.023, p = 0.160. **J.** Distribution of modulation indices from all DS, AS, BrT and SbC boutons during all trials (blue, N = 4839 boutons, median: −0.26) or during stationary trials only (orange, N = 4824 boutons, median = −0.219). Data from 8 FOV from 4 mice, using drifting gratings of 0.08 cpd. Red line indicates a modulation index value of zero (no change in response across arousal states). **K.** Cumulative distribution of modulation indices for data in J. 76.0% or 73.9% of boutons had negative modulation indices when either all trials or only stationary trials were included, respectively. Red line indicates a modulation index value of zero (no change in response across arousal states). **L.** Scatter plot of modulation indices of individual boutons, calculated from stationary periods or across all trials. Data from 8 FOV from 4 mice, using drifting gratings of 0.08 cpd. Red line: line of unity.

**Figure S3.**
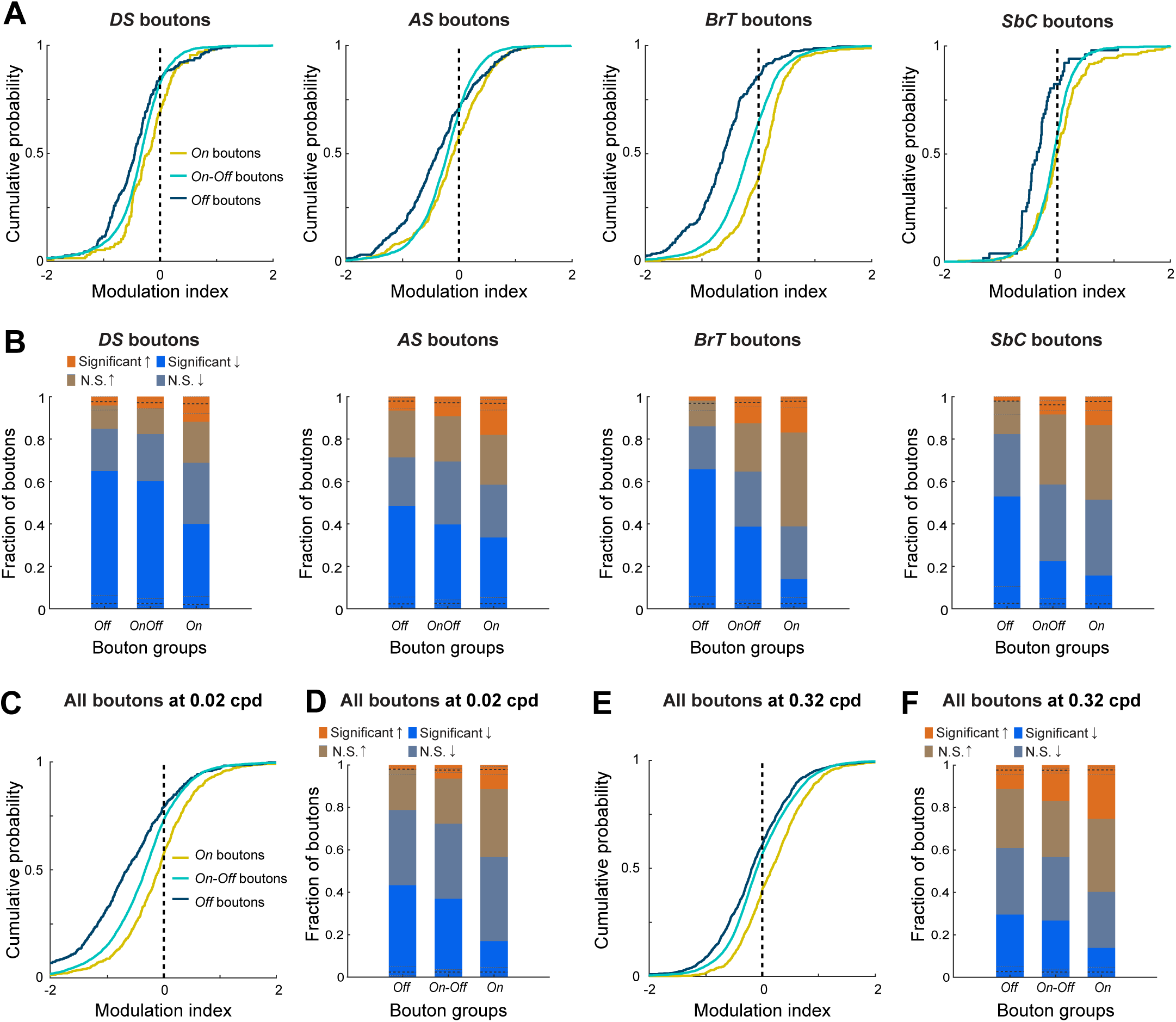
Responses of Off boutons are more suppressed by arousal regardless of spatial frequency, Related to Figure 3. **A.** Cumulative distribution of modulation indices of On, On-Off and Off boutons across broad functional categories. In each category, the distributions were significantly different between On and Off boutons, and between On-Off and Off boutons (p < 0.001, Kolmogorov-Smirnov test with Bonferroni correction for 3 comparisons). The distributions of On and On-Off boutons were also significantly different (p < 0.01, p < 0.001, 0 < 0.001, p=0.011 for DS, AS, BrT and SbC categories, Kolmogorov-Smirnov tests with Bonferroni correction). Numbers of On, On-Off and Off boutons in each category: DS: 135, 2822, 217; AS: 405, 2631, 276; BrT: 414, 2779, 272; SbC: 179, 1165, 51. Median modulation indices for On, On-Off and Off boutons in each category: DS: −0.202, −0.330, −0.446; AS: −0.097, −0.206, −0.392; BrT: 0.107, −0.183, −0.622; SbC: −0.012, −0.066, −0.361. The proportion of On, On-Off and Off boutons with negative indices: DS: 68.9%, 82.4%, 84.8%; AS: 58.5%, 69.4%, 71.4%; BrT: 38.9%, 64.7%, 86.0%; SbC: 51.4%, 58.6%, 82.4%. Indices were calculated from responses to drifting gratings at 0.08 cpd. **B.** Fraction of boutons per group (On, On-Off, or Off) that were significantly enhanced, not significantly (N.S.) enhanced, not significantly suppressed, or significantly suppressed. Black dashed lines indicate the fraction of boutons that were significantly enhanced or significantly suppressed in shuffled data (median across 1000 shuffles). Indices were calculated from responses to drifting gratings at 0.08 cpd. **C.** Cumulative distribution of modulation indices of On, On-Off, and Off boutons, calculated from responses to drifting gratings at 0.02 cpd. Distributions were significantly different (all p’s < 10-16, Kolmogorov-Smirnov tests with Bonferroni correction). Median indices for On, On-Off and Off boutons: −0.08, −0.32, and −0.64, respectively. 56.6%, 72.3%, and 78.8% of On, On-Off, and Off boutons had negative index values, respectively. Ns of On, On-Off and Off boutons were 732, 6278, and 462, respectively. **D.** Fraction of Off, On-Off, and On boutons that were significantly enhanced, not significantly (N.S.) enhanced, not significantly suppressed, or significantly suppressed. Black dashed lines indicate the fraction of boutons that were significantly enhanced or significantly suppressed in shuffled data (median across 1000 shuffles). Indices were calculated from responses to drifting gratings at 0.02 cpd. **E.** Cumulative distribution of modulation indices of On, On-Off, and Off boutons, calculated from responses to drifting gratings at 0.32 cpd. Distributions were significantly different (all p’s < 10-10, Kolmogorov-Smirnov tests with Bonferroni correction). Median indices for On, On-Off and Off boutons are 0.173, −0.095, −0.185, respectively. 40.3%, 56.7%, 60.9% of On, On-Off, and Off boutons had negative index values, respectively. Ns of On, On-Off and Off boutons were 976, 8173, 1280. **F.** Same as D, but with indices calculated from responses to drifting gratings at 0.32 cpd.

**Figure S4.**
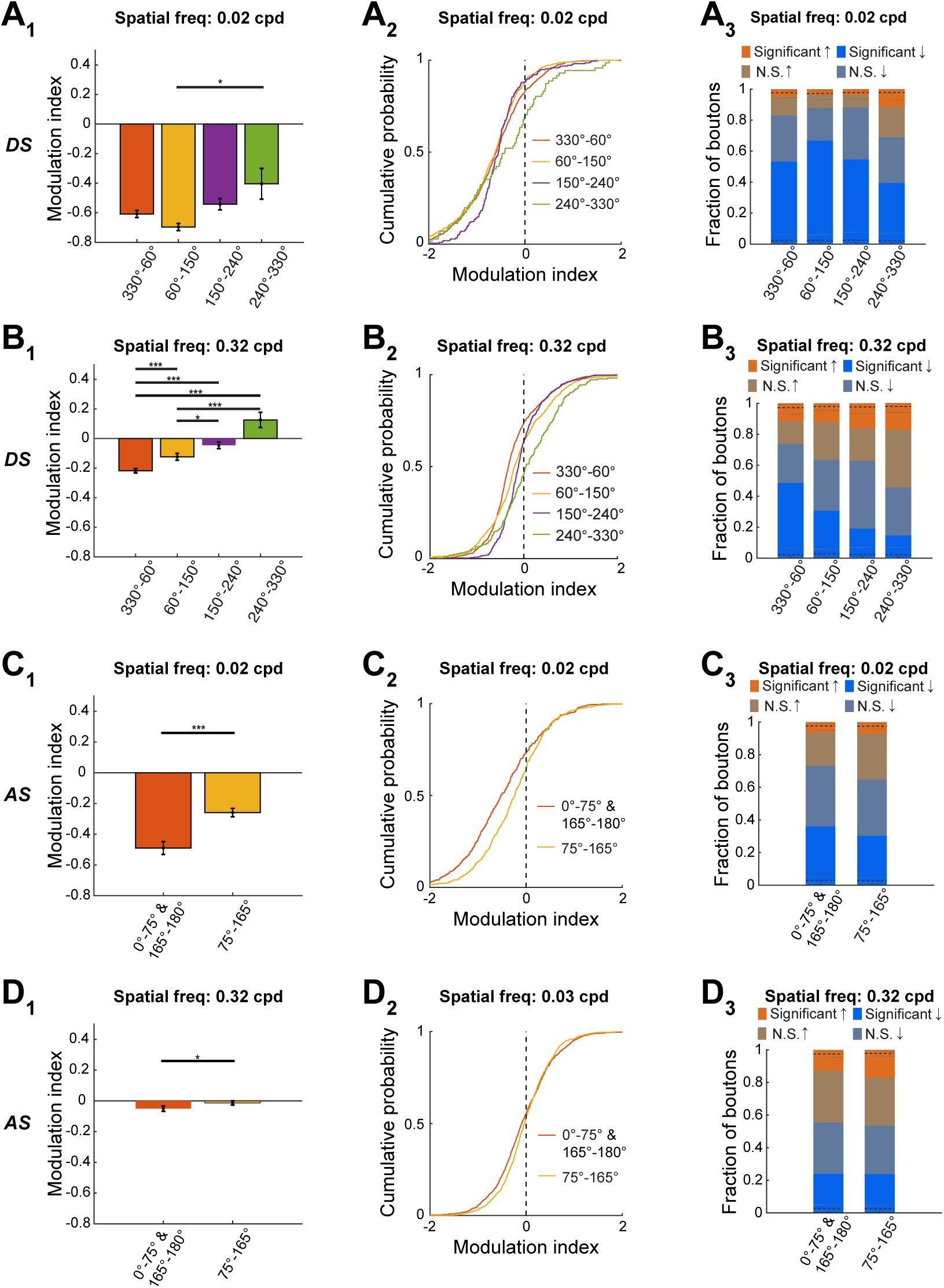
Arousal differentially modulates boutons with distinct motion preferences or receptive field locations, Related to Figure 4. **A. A_1_**, average arousal modulation index of DS boutons in each group. Mean ± SEM. Median values for groups of boutons with preferences ranging from 330°-60°, 60°-150°, 150°-240°, 240°-330° were −0.600, −0.620, −0.554, −0.426, respectively. Distributions of indices were significantly different across groups (p = 0.011, Kruskal-Wallis test with Dunn correction; *p < 0.05). Ns for each group: 852, 796, 214, 71. **A_2_**, cumulative distributions of modulation indices for each group. Indices were negative for 83.1%, 87.9%, 88.3%, 69.0% of boutons in each group. The distribution of the 330°-60° group was not significantly different from the 60°-150° group (p = 0.363) but was significantly different from the 150°-240° group (p = 0.014), and the 240°-330° group (p = 0.037). The distribution of the 60°-150° group was significantly different from the 150°-240° group (p < 0.01), and the 240°-330° group (p < 0.001). The distribution of the 150°-240° group was significantly different from the 240°-330° group (p < 0.01). Kolmogorov-Smirnov tests with Bonferroni correction. A_3_, Fraction of boutons in each group that were significantly enhanced, not significantly (N.S.) enhanced, not significantly suppressed, or significantly suppressed. Black dashed lines indicate the fraction of boutons that were significantly enhanced or significantly suppressed in shuffled data (median across 1000 shuffles). Indices were calculated from responses to drifting gratings at 0.02 cpd. **B.** The same as A, but for presentation of gratings at 0.32 cpd. **B_1_**, mean ± SEM. Median value for groups boutons with preferences ranging from 330°-60°, 60°-150°, 150°-240°, 240°-330° were: −0.326, −0.197, −0.112, 0.047, respectively. Distributions of indices were significantly different across groups (p < 10-20, Kruskal-Wallis test with Dunn correction; *p < 0.05, ***p<0.001). Ns for each group: 1454, 910, 419, 218. **B_2_**, indices were negative for 73.9%, 63.5%, 63.0%, and 45.9% of boutons in each group. Distributions were significantly different between all the groups (all p’s < 10-4, Kolmogorov-Smirnov test with Bonferroni correction). **B_3_**, Same as **A_3_**, but with indices calculated from responses to drifting gratings at 0.32 cpd. **C. C_1_**, average arousal modulation index of AS boutons in each group. Mean ± SEM. Median values for groups of boutons with preferences ranging from 345°-75° and 75°-165° were −0.527 and −0.236, respectively. Distributions of indices were significantly different across groups (***p < 10-5, Wilcoxon rank sum test). Ns for each group: 388, 675. **C_2_**, cumulative distributions of modulation indices for each group. Indices were negative for 73.2% and 64.9% of boutons in the 0°-75° & 165°-180° and 75°-165° groups, respectively. Distributions were significantly different (p < 10-5, Kolmogorov-Smirnov test). **C_3_**, Fraction of boutons in each group that were significantly enhanced, not significantly (N.S.) enhanced, not significantly suppressed, or significantly suppressed. Black dashed lines indicate the fraction of boutons that were significantly enhanced or significantly suppressed in shuffled data (median across 1000 shuffles). Indices were calculated from responses to drifting gratings at 0.02 cpd. **D.** The same as **C**, but for presentation of gratings at 0.32 cpd. **D_1_**, mean ± SEM. Median values for groups of boutons with preferences ranging from 0°-75° & 165°-180° and 75°-165° were −0.092, −0.038, respectively. Distributions of indices were significantly different across groups (*p = 0.024, Wilcoxon rank sum test). Ns for each: 1384, 1495. **D_2_**, indices were negative for 55.6%, 53.6% of boutons in each group. The two distributions were significantly different (p < 10-3, Kolmogorov-Smirnov test). **D_3_**, Same as **C_3_**, but with indices calculated from responses to drifting gratings at 0.32 cpd.

**Figure S5.**
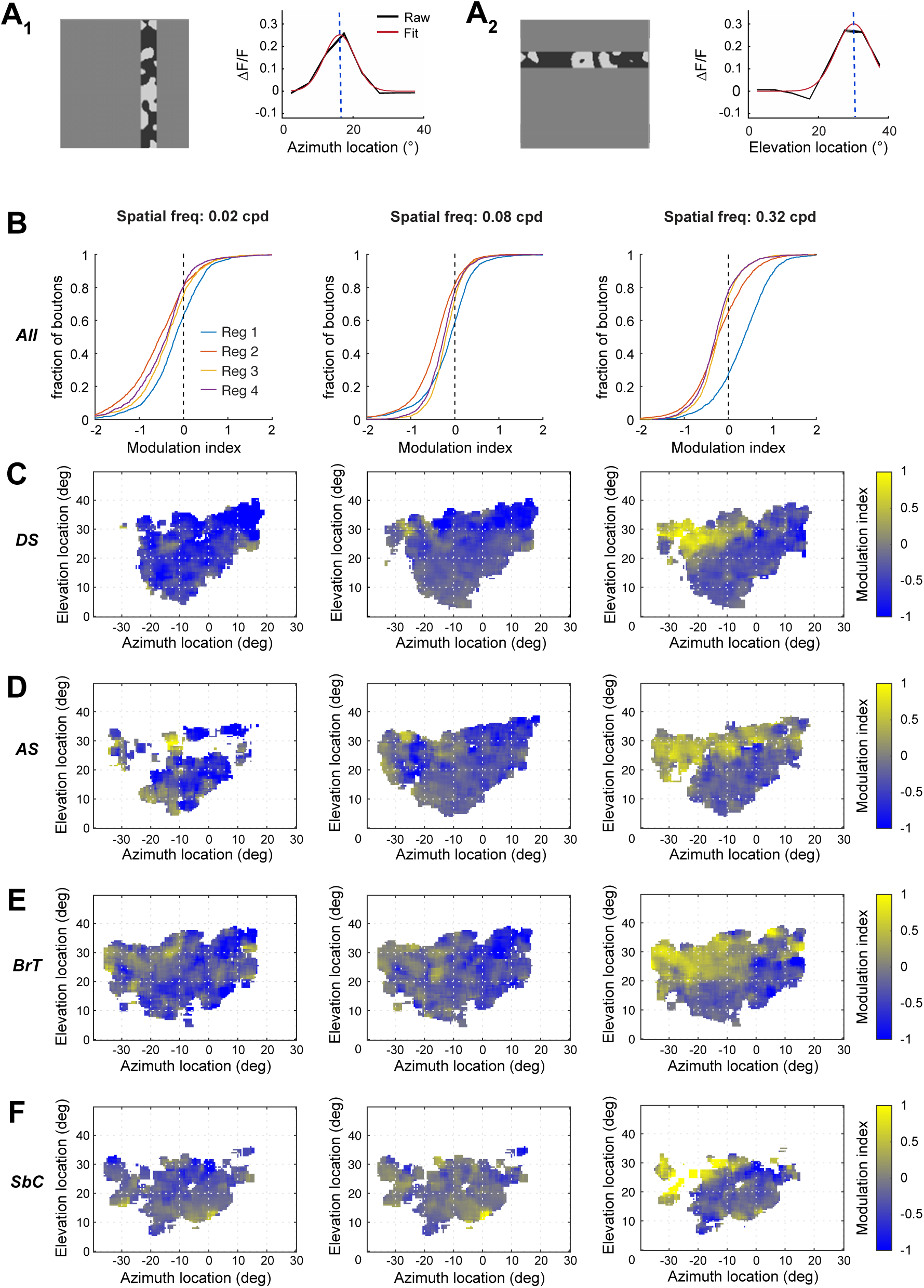
Arousal-related response suppression or enhancement in retinal boutons depends on retinotopic preference, Related to Figure 4. **A.** Response curves of two example boutons to a 5° x 40° bar containing spatiotemporal noise displayed at one of 8 azimuth locations (**A_1_**) or one of 8 elevation locations (**A_2_**) on the monitor. **B.** Cumulative distributions of modulation indices for all the boutons (DS, AS, BrT and SbC) in each region (defined by the boutons’ retinotopic preferences, see Figure 4C_2_), for stimuli with one of three spatial frequencies. The distributions between any two regions were significantly different (all p’s < 0.001), except for between Region 3 and Region 4 at 0.02 cpd (p = 0.08) and at 0.32 cpd (p < 0.01). Kolmogorov-Smirnov tests with Bonferroni correction. **C.** Modulation indices of individual direction-selective (DS) boutons mapped to their preferred retinotopic locations in the visual space, when using responses to drifting gratings with spatial frequencies of 0.02 cpd, 0.08 cpd or 0.32 cpd. **D.** Same as C but for modulation indices of individual axis-selective (AS) boutons. **E.** Same as C but for modulation indices of individual broadly-tuned (BrT) boutons. **F.** Same as C but for modulation indices of individual suppressed-by-contrast (SbC) boutons.

## STAR METHODS

### CONTACT FOR REAGENT AND RESOURCE SHARING

Further information and requests for resources and reagents should be directed to, and will be fulfilled by, the Lead Contact, Mark L. Andermann.

### EXPERIMENTAL MODEL AND SUBJECT DETAILS

All animal care and experimental procedures were approved by the Beth Israel Deaconess Medical Center Institutional Animal Care and Use Committee. Animals were housed with standard mouse chow and water provided *ad libitum*. Male C57BL/6 adult mice (2-6 months old) were used in this study.

### METHOD DETAILS

#### Viral injection

To label retinal ganglion cell axons, 1.2 µl of AAV2/2.CAG.GCaMP6f.WPRE.SV40 ([1]; Boston Children’s Hospital Viral Core) was gently injected intravitreally into the right eye after the mouse was anesthetized using isoflurane in 100% O_2_ (induction, 3%–5%; maintenance, 1%– 2%). Care was taken to minimize bleeding and to prevent cataract formation during the injection procedure.

#### Headpost and cranial window implantation

A headpost and cranial window were implanted 2-3 weeks after viral injection. Mice were given 0.03 ml of dexamethasone sodium phosphate (4 mg/ml, i.m.) roughly 3 hours prior to surgery in order to reduce brain edema. Mice were anesthetized using isoflurane in 100% O_2_ (induction, 3%–5%; maintenance, 1%–2%) and placed on a heating pad (CWE) in a stereotaxic apparatus (KOPF). Ophthalmic ointment (Vetropolycin) was applied to the eyes. Using a procedure similar to that described previously [2], a two-pronged headpost was affixed to the skull, centered roughly 2.7 mm lateral and 1.9 mm posterior to Bregma over the left hemisphere, tangential to the curved skull surface. The head was then tilted to secure the headpost in custom clamps (Thorlabs, Standa) that aligned the headpost precisely parallel to the platform of the stereotaxic apparatus. A 3-mm diameter craniotomy was performed at the center of the headpost. The underlying cortical and hippocampal tissue was carefully aspirated until reaching the surface of the thalamus. The thalamic surface and optic tract were kept intact. A 3-mm diameter coverslip (glued to the bottom of a 3 mm x 3.4 mm (diameter x height) stainless steel cylindrical cannula (MicroGroup) prior to surgery using UV-cured Norland Optical Adhesive 71) was lowered into the craniotomy, approximately 2.75 mm below the skull, where it pressed slightly on the surface of the thalamus. The cannula was affixed to the skull with Vetbond (3M) followed by C&B Metabond (Parkell), to form a permanent seal. To create a low-profile adaptor to accommodate the water-immersion objective and light shielding, a neodymium ring magnet (Indigo® Instruments, outer diameter, inner diameter, height: 7.5 mm, 5 mm, 1 mm) was positioned around the cannula and glued to the skull. During two-photon imaging sessions, this ring magnet held the light shielding in place by contact with a 8 mm x 0.3 mm (diameter x height) spring steel round shim (McMaster) attached to the blackout fabric (Thorlabs). Meloxicam (0.5 mg/kg, s.c.) was administered and the mouse was allowed to recover.

#### Epifluorescence and two-photon calcium imaging

To measure bulk calcium signals in response to visual stimulation in head-fixed mice across arousal states, we performed epifluorescence calcium imaging (Neurolabware). A blue LED light source (470 nm center, 40 nm band, Chroma) was used for excitation, and the green fluorescence was passed through a 500 nm long-pass emission filter and collected using a sCMOS camera (PCO). Light shielding around the objective was used to block light emitted by the LCD screen. We focused on the surface of the thalamus. Epifluorescence images (512 x 384 pixels) were acquired at 15.5 Hz, using a 4x, 0.28 NA objective (Olympus).

Two-photon calcium imaging was performed using a resonant-scanning two-photon microscope (Neurolabware). All images were acquired using a 20x, 1.0 NA, 5.6 mm WD objective (Zeiss) at 4.7x (∼160 x 210 µm^2^) or 2.8x (∼95 x 125 µm^2^) digital zoom. Light shielding around the objective was used to block light emitted from the LCD screen. We concentrated on imaging fields of view (FOV) at depths of 80-150 µm below the surface of the optic tract (roughly corresponding to the upper 20-90 µm of the dLGN shell; high-quality images could be obtained throughout the upper ∼140-150 µm of the dLGN, data not shown), using a MaiTai HP DeepSee Ti:Sapphire laser (80 MHz; Spectra-Physics) or an Insight X3 laser (80 MHz; Spectra-Physics) at 960 nm. The arousal modulation results were not sensitive to imaging depth within this range (not shown). Laser power measured at the front aperture of the objective was 30-65 mW – likely a substantial overestimate of actual power reaching the sample via the cannula. Images were collected at 15.5 frames/s, 686 × 512 pixels/frame, using Scanbox (Neurolabware). Each imaging run lasted approximately 30 min, and 4-5 runs were performed during each imaging session. Occasionally, the imaging depth was adjusted slightly in between runs to account for slow drifts in the z-plane. For a given mouse, each FOV imaged in a given session was at least 20 µm above or below any FOV imaged in another session. Epifluorescence and two-photon imaging experiments were typically performed between one week and one month after headpost and cranial window implantation.

#### Pupil videography and image analysis

During epifluorescence imaging, the eye contralateral to visual stimulation was illuminated by an infrared LED. During two-photon imaging, the pupil was illuminated by the spread of two-photon excitation infrared light emanating from within the brain. Images of a pre-selected region of interest around the eye were recorded using a CCD camera (Dalsa) at 15.5 Hz, synchronized with fluorescence image acquisition (Scanbox, Neurolabware).

#### Locomotion during two-photon imaging

Locomotion was recorded using a customized rotary encoder at a resolution of 0.5 cm/step. Total distance travelled was registered at 15.5 Hz, synchronized with two-photon image acquisition (Scanbox, Neurolabware). The running speed was calculated as the displacement between two adjacent data points dividing by the time between data points (∼.065 s), and smoothed by a median filter (width: 5 samples, or ∼.32 s).

#### Visual Stimulation

Visual stimuli were generated using Psychtoolbox (Brainard, 1997), and displayed on a luminance-calibrated LCD monitor (Dell, 17”,1280 x 1024 pixels, 60 Hz refresh rate) placed 22 cm from the mouse’s right eye and spanning 80° × 70° of visual space (azimuth: −40° to 40°, with 0° indicating the line perpendicular to the eye, which is ∼45° lateral to the axis in front of the head, see Figure 4C_1_; elevation: −13° – 57° relative to this same line).

To measure arousal modulation during epifluorescence imaging, we presented full-screen sine-wave drifting gratings (80% contrast) at 0° (nasal to temporal direction) and at spatial frequencies of 0.02, 0.08 and 0.32 cycles per degree (cpd) and a temporal frequency of 2 Hz. Gratings of different spatial frequencies were delivered in a random order, with blank trials (full-screen mean luminance – gray screen) interleaved in between. All stimuli were displayed for a 2-second duration. The inter-stimulus interval (mean luminance gray) lasted 2 seconds. One run per session consisted of 216 repeats of gratings at each spatial frequency and 54 blank trials. Recordings from the same animal were repeated on two days, and modulation indices from the two days were averaged to give the modulation index for the animal.

To measure arousal modulation during two-photon imaging, we presented full-screen sine-wave drifting gratings (80% contrast) at one of eight directions of motion spaced 45° apart, at spatial frequencies of 0.02, 0.08 and 0.32 cycles per degree and a temporal frequency of 2 Hz. The visual stimulation paradigm also included periods of full-screen mean luminance (gray, blank trials) or periods of luminance increments or decrements (‘On’ or ‘Off’ trials, respectively, 80% contrast). All stimuli were displayed for a 2-second duration. The inter-stimulus interval (mean luminance gray) lasted 2 seconds. A single repeat involved presentation, in random order, of the set of all of the above stimuli (one presentation of each direction and spatial frequency, three presentations each of ‘On’ and ‘Off’, and three or six presentations of ‘blank’ stimuli). A single run usually consisted of 10-14 repeats, each with a different randomization of stimulus order. We recorded 3-5 runs for each FOV in a given imaging session.

To measure retinotopy with high spatial resolution during two-photon imaging, we used a binarized version of a bandpass-filtered noise stimulus with a spatial frequency corner of 0.05 cycles per degree, a cutoff of 0.32 cycles per degree and a temporal frequency cutoff of 4 Hz [3]. The noise stimulus was presented within 5° × 40° bars, presented vertically at one of 8 azimuth locations and horizontally at one of 8 elevations. Stimuli were presented for 2 seconds each, with a 2-second inter-stimulus interval (mean luminance). Visual stimulation also included a blank condition (mean luminance). Stimulus order was randomized within a single repeat (consisting of a single presentation of each stimulus condition), and 30 repeats were presented during one run. We performed 1-2 runs of retinotopic mapping per imaging session.

#### Image processing

##### Image preprocessing

To correct for x-y motion along the imaged plane, a series of image registration and data cleaning steps were applied [2]. For epifluorescence images, movies collected on each imaging day were registered to a common average field-of-view using a local image normalization method (subtraction of local mean and division by local variance across pixels) and an image warping method (imregdemons.m, Matlab, Mathworks). For two-photon images, additional denoising method was implemented to aid image registration. Specifically, principal component analysis (PCA) was performed to identify spatial principal components of the field of view and each image of the movie was reconstructed using only the first 400 (highest eigenvalues) out of ∼30,000 principle components. The denoised images were then subjected to local normalization and image warping to derive the warping parameters, which were then applied to the original raw movie. For additional details, please see [2].

##### Identification of the responsive region in the dLGN from bulk calcium imaging

The dLGN region could be easily distinguished from superior colliculus or other thalamic regions, based on baseline green fluorescence from GCaMP6f-expressing retinal ganglion cell axons, and from the characteristic oval shape. However, given the limited dimensions of the LCD monitor used for displaying visual stimuli, and the obstructions in the upper visual field due to light shielding, not all the dLGN regions receiving RGC projection responded to visual stimulation. For quantifying calcium signals across arousal states, we only considered the portion of the dLGN that showed robust visually evoked responses during at least one of the stimulus conditions (spatial frequencies). The visually evoked response of a given pixel in the acquired image was defined as robustly driven if the average responses was two standard deviations above the average activity estimated during blank trials.

##### Identification of bouton masks

To identify boutons and extract masks for further signal processing, we established an automated image segmentation algorithm. First, an image of the absolute value of the average ΔF/F was calculated for each trial type by averaging single condition evoked response maps across all N repetitions of that trial type (|mean((F_i_-F_i0_)/F_i0_)|, i=1..N, where F_i_ is the mean fluorescence during the stimulus presentation and F_i0_ is the average baseline fluorescence during the 1 second prior to each stimulus onset). We used the absolute value of ΔF/F in order to include boutons that were strongly suppressed by visual stimuli, corresponding to negative values of ΔF/F. A bouton identification procedure was then independently applied to each of these average ΔF/F images. First, local normalization was applied, with the local mean estimated by isotropic filtering of the image using a Gaussian kernel (with standard deviation, sigma = 3 µm). The local variance was estimated using a larger Gaussian filter (sigma = 50 µm).

Morphological filters were then applied to identify, in an unbiased manner, connected sets of pixels that together resembled the size and shape of a typical RGC bouton, as follows: first, small pixel gaps were filled by interpolation using a square-shaped structuring element of 1.3 x 1.3 µm^2^. We then removed all small unconnected structures via an ‘opening’ operation using the same structuring element. To obtain candidate masks, we first binarized the above images by setting to ‘1’ all pixels with values above 10% to 15% of the maximal pixel amplitude after filtering, and setting all other pixels to ‘0’. A Euclidian distance transform was then applied to these binary images (MATLAB function ‘bwdist.m’). The built-in MATLAB watershed transform was used to finalize the segmentation. The results from the distance transform and the watershed transform from the individual ΔF/F images were combined by summing the distance transform across conditions and normalizing this value by the bouton count obtained by the watershed transformation. A final watershed transformation was applied to this normalized distance image to increase the accuracy of the procedure and to reduce the risk of false positives in the bouton identification procedure. In addition, to remove residual calcium signals not originating from the bouton itself, we estimated neuropil masks as circular annuli of 3 µm width, with the inner edge located 2 µm beyond the edge of a corresponding bouton mask. Pixels from adjacent bouton masks were excluded from these neuropil masks.

##### Timecourse extraction and correction

To obtain raw fluorescence traces for dLGN region in epifluorescence imaging datasets, or for bouton masks and neuropil masks in two-photon imaging datasets, the fluorescence intensity value at each time point was defined as the average fluorescence across the pixels belonging to the region or to the mask.

To account for photobleaching during two-photon imaging sessions, a bleaching correction method was established. Raw bouton and neuropil traces were first smoothed using a sliding filter (30^th^ percentile of a 5-minute sliding window). Then, the filtered traces were fitted using a decaying exponential, where the amplitude and the offset were independently estimated for each bouton and each neuropil ring, while the time constant was fixed to an experimentally defined constant value of 75 minutes, which was in agreement with time constants other groups have determined for the photobleaching of GCaMP6f using two-photon imaging at similar excitation wavelengths and laser power (Harris lab/Photophysics, https://www.janelia.org/lab/harris-lab-apig/research/photophysics/two-photon-fluorescent-probes). To correct for photobleaching in each trace, the fitted offset value was first subtracted from the raw trace, and then the resulting trace was multiplied by the inverse of the exponential of the fixed decay time constant before adding back the offset value.

To account for neuropil signals which may contaminate signals in the bouton trace, neuropil correction was applied by subtracting a scaled version of the corresponding neuropil trace (0.6 x neuropil trace) from each bouton trace before adding back the mean neuropil fluorescence (temporally-averaged across the neuropil trace) [4]. (Kerlin et al., 2010).

To account for autofluorescence from brain tissue in the epifluorescence image, we estimated the average fluorescence value from a region anterior to the dLGN which received no RGC input, which served as approximation of the level of background autofluorescence. We subtracted this estimate on a frame-by-frame basis from the fluorescence trace of the identified dLGN region.

We also corrected baseline fluorescence, F_0_, to remove the decay in fluorescence from activity evoked during the previous visual stimulus presentation. Due to slow decay dynamics *in vivo* (as a result of calcium buffering and GCaMP6f buffering) after stimulus-evoked calcium activity, the bouton fluorescence sometimes did not fully return to baseline during the 1 second after the offset of the previous stimulus presentation, and persisted in the 1 second used to calculate F_0_ for the following stimulus period. Therefore, a baseline correction was introduced that modeled this exponential decay of previously evoked GCaMP6f calcium activity, using an experimentally determined fixed time constant of 1 second (in agreement with previously determined GCaMP6f dynamics *in vivo*; [3]).

To assess the fractional change in fluorescence, ΔF/F(t), during each visual stimulus presentation, the fitted, exponentially decaying contribution from the previous trial was first subtracted from F(t) during the 1-second interval prior to and the 2-second interval during visual stimulus presentation. Then, the corrected baseline was used as the new baseline F_0_ to compute ΔF/F(t). Single response values during a given trial were obtained by averaging the ΔF/F(t) response during the 2-second stimulus window.

#### Estimation of visual tuning preferences in RGC boutons

##### Estimation of boutons with significant visual responses

We define single condition bouton responses in two-photon imaging datasets as the average from all the trials of a given stimulus condition. Visually evoked responses were corrected by subtracting the average response across blank trials. There were approximately 30 repeats for each of the retinotopic conditions, 60 trials for each combination of direction and spatial frequency (SF), 180 trials of luminance increments and decrements (‘On’ and ‘Off’) and 360 blank trials. Different boutons had differing response dynamics, and we attempted to minimize bias in which boutons were deemed significantly visually responsive. Thus, we assessed, for each bouton and each stimulus condition, if the evoked response was significantly different from noise, by requiring the amplitude of ΔF/F(t) during the response window to exceed 2.5 standard deviations above or below the mean baseline activity (computed using the 1-second window prior to stimulus onset) for at least 10 out of the 31 time points (15.5 Hz frame rate x 2 sec stimulus presentation). For assessment of significant On and Off responses, we only required the ΔF/F(t) amplitude to exceed this threshold for at least 5 of the 31 time points, as a substantial proportion of boutons exhibited transient On and/or Off responses. These criteria were confirmed to be highly conservative, thereby including only highly robustly visually responsive boutons.

To assess if a given bouton exhibited *a significantly positive response* at a particular spatial frequency, we required that at least 3 out of the 8 directions at this spatial frequency evoked significantly positive responses according to the criteria described above. A similar approach was used to determine if a bouton exhibited a significantly negative response to a particular spatial frequency. Note that all boutons contributing to the main results underwent additional quality controls (see below), further protecting against inclusion of any noisy boutons in our analyses.

##### Direction tuning curve fitting

For each bouton showing a significantly positive response at a given spatial frequency, a direction tuning curve was computed across directions for stimuli presented at that spatial frequency. The direction tuning curves were initially sampled in steps of 45°. In order to obtain a more precise estimate of the preferred axis and direction, a fitting approach was used to estimate the preferred direction. Tuning curves were fitted with a two-peaked Gaussian with offset [5]:

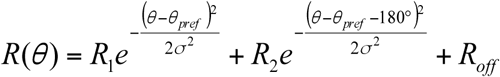

*R*(*θ*) was the ΔF/F response for stimulus direction *θ*. This model assumed that the peaks of the two Gaussians were 180° apart. *θ_pref_* was defined as the preferred direction evoking the strongest ΔF/F response, *R*_1_. *R*_2_ was the amplitude of the second peak located at *θ_pref_* + 180°. It was also assumed that both Gaussians shared a common standard deviation, *σ*. The fifth fitted parameter was a constant amplitude offset, *R_off_*.

Several steps were taken to improve the reliability of the fitting of direction tuning curves and to optimize the accuracy of estimation of preferred direction of motion. To increase the number of input points for the fitting procedure from 8 to 25, a heuristic method of interpolation and extrapolation was implemented. First, a ninth point was added at 360°, which was identical to the one at 0°. Then, the number of input points was doubled from 9 to 17 by linear interpolation of the 9-point direction tuning curve. For the interpolated data point between the two most strongly driven initial directions (out of 9), we further adjusted the interpolated amplitude so that its value became a close approximation of that predicted point from a Gaussian curve fit through the rest of the points, thus reducing the error introduced by linear interpolation given the expected continuity of the curves. To this end, we applied the empirical formula described below. Note that our results were largely unchanged if this additional adjustment to the linear interpolation was omitted (mean difference in preferred direction: 2°, median difference: 1.2°). However, this additional peak adjustment resulted in significantly smaller residual values between the fitted curve and the initial 8-direction tuning curve.

The interpolated amplitude between the two most strongly driven initial directions was calculated as follows. *R_S_*_1_ was defined as the strongest response out of all 8 directions and *R_S_*_2_ as the stronger of the two responses for directions ± 45° adjacent to *R_S_*_1_. *R_S_* _3_ was defined as the weaker of the two responses adjacent to *R_S_*_1_. *R_S_* _4_ was defined as the response adjacent to *R_S_* _2_, at 90° from *R_S_*_1_. The interpolated ΔF/F response *R_S_*_12_ between *R_S_*_1_ and *R_S_* _2_ was defined as: 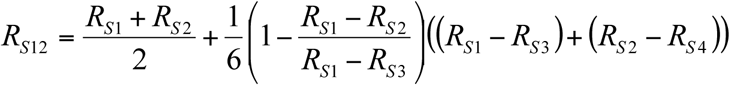. This method compared the slope between *R_S_*_1_ and *R_S_* _2_ with the slope between *R_S_*_1_and *R_S_* _3_. If the peak was flat, a maximum amount of 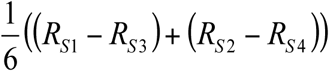 was added, corresponding roughly to the expected value of a Gaussian peak. If the absolute values of the slopes between *R_S_*_1_ and *R_S_* _2_ and between *R_S_*_1_ and *R_S_* _3_ were identical (and therefore *R_S_*_1_ was the real peak of the Gaussian), this corresponds to 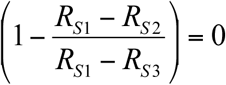 in the above equation, therefore resulting in no additional value added to the interpolation method. A similar method was used to interpolate negative peaks.

To further improve the stability of the fitting procedure and to better approximate the direction tuning curve, we added two shadow-copies of the two-peaked Gaussian function, circularly shifted by +360° and −360°:

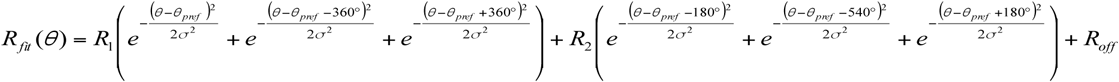

This addition of the shadow-copies increased the range to [-90°, 450°], and thus extended the fitted tuning curve by 4 additional directions (at 22.5° spacing) on either end. While the adjusted linear interpolation and the addition of the shadow copies improved the fitting procedure, similar results were obtained using the basic 17-point linearly interpolated tuning curve (data not shown).

A bootstrapping method involving random sampling of trials from each condition was then implemented to fit the tuning curves. Specifically, for each of 100 iterations, the tuning curve was initially computed by randomly sampling with replacement, and by averaging responses from 60 trials sampled from each of the 8 directions. These 8-point tuning curves were then interpolated, extended and finally fitted using the method described above. The final parameters used were the mean of the fitted parameters across the 100 sampling iterations.

To determine if the fitting procedure yielded a high-quality fit, a combination of criteria was used. Each iteration of the fitting procedure yielded a coefficient of determination, *r*^2^, defined as the explained variance using least-squares regression to fit the data. As a second control step, a combined coefficient of determination, 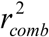, was computed by comparing the original direction tuning curve with the fitted curve derived using the average of each fitting parameter (across 100 iterations). To assess both the quality and the reliability of the fitting procedure, we introduced a heuristic goodness of fit, *G_fit_*:

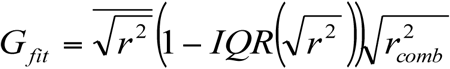 where *IQR* was defined as the interquartile range – the difference between the 75^th^-percentile and the 25^th^-percentile (of 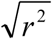 values across iterations). A bouton was considered to have a well-fit direction tuning curve at a given spatial frequency if the goodness of fit, *G*, was greater than 0.66. The threshold was chosen based on examination of a large proportion of example boutons, and values in the range of 0.5 to 0.9 yielded similar results. The complete direction curve fitting procedure was separately run for each of the three spatial frequencies employed, and therefore each bouton was attributed up to three sets of fitting parameters.

##### Axis and direction selectivity

For each bouton, we calculated its axis and/or direction selectivity at a given spatial frequency if we observed a significantly positive evoked response at that spatial frequency. For axis selectivity, we calculated a ‘vector sum’ axis selectivity index (ASI; i.e., selectivity for a motion along a given axis) on each interpolated direction tuning curve [4]. This index was calculated by projecting the ΔF/F response for each of the 16 directions in the range between 0° and 360° onto a circle with 2*i* progression and estimating the magnitude of the normalized vector sum, which ranged from 0 to 1 (maximum selectivity): 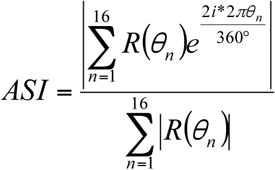. Opposite directions contributed in an additive fashion, while orthogonal directions canceled each other out. The ASI computation was iterated 100 times by bootstrapping and averaged for each spatial frequency.

In a similar manner, we computed a ‘vector sum’ direction selectivity index (DSI), by projecting the 16 directions onto a circle with 1*i* progression: 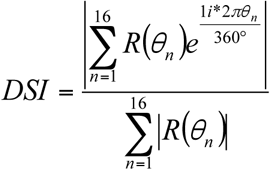. As with ASI, the DSI estimate was repeated with the bootstrapping method, and the average was used as the DSI for that spatial frequency.

##### Preferred direction of motion and preferred axis of motion

For direction-selective boutons or axis-selective boutons, the preferred direction of motion or preferred axis of motion was defined as the direction or axis of the mean vector across the preferred motion direction or motion axis for each spatial frequency, which was the direction or axis with peak response amplitude estimated from the fitted tuning curve at the given spatial frequency. Estimates of preferred direction or axis of motion ranged from 0° and 360°, or 0° and 180°, respectively.

To compensate for the pitch angle of the mouse head during imaging acquisition (Pitch of 31.6° above the angle estimated for a typical ambulatory position [6]; Roll: 16.7°; measured using stereotaxic coordinates during headpost implant, N=3 mice), we rotated the preferred direction or axis of motion counterclockwise by 31.6°.

#### Bouton type classification

A bouton was classified as direction selective if, for at least one of the three spatial frequencies used, (i) it had a significant positive response, (ii) the tuning curve was successfully fit (as estimated by goodness of fit criteria), (iii) the direction selectivity index (DSI) exceeded 0.2 (a value equivalent to 0.33 if direction selectivity was calculated as 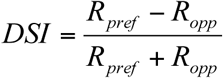, where *R_pref_* was the response at the preferred direction and *R_opp_* was the response at the opposite direction). Additionally, for all spatial frequencies which showed a significant positive response, we also required the DSI at each of these spatial frequencies to exceed 0.15 (to ensure that group assignment was not sensitive to the vagaries of which spatial frequencies were used). Finally, we verified that boutons in this group did not show any significantly negative responses for any spatial frequency.

A bouton was defined as axis selective (i.e., most strongly responsive to motion along two opposite directions constituting a single axis of motion) if (i) it had at least one significantly positive response for at least one of the three spatial frequencies, (ii) the fitting procedure was reliable, and (iii) the axis selectivity index (ASI) exceeded 0.15 (a value equivalent to 0.33 if axis selectivity was calculated as 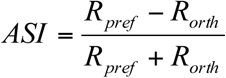, where *R_pref_* was the response at the preferred direction and *R_orth_* was the mean response at the two directions orthogonal to the preferred one). We distinguish the term axis selectivity from orientation selectivity, as the latter is often used even for responses to stationary (i.e., non-drifting) oriented gratings – a stimulus not examined in this study. It is possible that certain axis selective boutons may not be strongly driven by stationary gratings. For all spatial frequencies for which we observed a significant positive response, we further required that the DSI was less than 0.2 and that the ASI was greater than 0.1. If these conditions were not met or if the bouton showed any significant negative response at any spatial frequency, the bouton was removed from this group.

A bouton was defined as ‘broadly tuned’ if (i) it exhibited a significant positive response for at least one of the three spatial frequencies and (ii) the ASI and DSI were below 0.15 and 0.2, respectively, for all significant spatial frequencies. We also removed boutons that exhibited any significant negative responses at any spatial frequency.

Boutons having significant negative responses for at least one spatial frequency and no significant positive response at any spatial frequency were defined as ‘suppressed’. ‘Suppressed-by-contrast’ (SbC) boutons were further defined as those boutons generally suppressed by all types of visual stimuli, including horizontal and vertical bars containing spatiotemporal noise (used for retinotopic mapping). Moreover, a SbC bouton needed to be significantly suppressed by at least 2 out of the 8 retinotopic conditions in at least one of the two stimulation axes. If no retinotopic mapping stimulus evoked a significant response, or if the response to one or both retinotopic stimulation axes was significantly positive, the bouton was classified as ‘suppressed’, but not as part of the subcategory of ‘SbC’ boutons. Only the SbC subcategory of suppressed boutons was considered in this study.

Boutons which showed a significant visually-evoked response but were not classified into any of the above conservatively-defined categories, were ‘unclassified’ boutons and were not included in subsequent analyses.

Finally, a small proportion of candidate bouton masks were not significantly driven by any of the presented visual stimuli. These were classified as ‘unresponsive’ and not considered further.

#### Arousal modulation

The arousal level of the mouse was assessed by estimating pupil area [7–11]. Pupil area was automatically identified from the acquired videos using customized scripts in Matlab (MathWorks). For extracting the pupil diameter when recorded during two-photon imaging, the Starburst algorithm was used [12]. For extracting the pupil diameter when recorded during epifluorescence imaging, morphological filters and the grayconnected.m function (Matlab, MathWorks) were first used to identify the region of image that belonged to the pupil. The regionprops.m function (Matlab, Mathworks) was then used to fit an ellipse to the pupil region and to measure parameters of the ellipse. Pupil area was calculated as the area of the fitted ellipse and smoothed by a median filter (width: 5 adjacent frames).

The pupil area corresponding to each image frame was first calculated from the fitted ellipse (see above). The distribution of pupil areas across all runs on the same day was then assessed, and the area values at 25% and 75% were selected to be the thresholds for low arousal and high arousal states. For each trial, the pupil area estimates from 1 sec before to 2 sec after onset of visual stimulation were averaged, and the result was used as the pupil area for that trial. If the average pupil area for a given trial was below the 25% threshold, the mouse was considered to be in a low-arousal state during that trial. If the average pupil area was above the 75% threshold, the mouse was considered to be in a high-arousal state.

To calculate the correlation between visual responses and pupil area, the average visually evoked ΔF/F during the 2 seconds of visual stimulation was used as the response for that trial. The Pearson correlation method was used to quantify the correlation between visually evoked ΔF/F and pupil area across all presentations of a given stimulus.

To quantify the degree of arousal modulation of responses of drifting gratings, we defined a modulation index, MI= [R_HighArousal_ – R_LowArousal_]/R_AllTrials_, where R_HighArousal_, R_LowArousal_, R_AllTrial_ represent the average response across all the high arousal trials, the average response across all the low arousal trials, and the average response across all trials, respectively. For epifluorescence imaging, these were responses to 0° drifting gratings. For individual boutons from two-photon imaging, we considered three directions: the direction that drove the maximal response, and the two adjacent directions. We first calculated the average response across trials for each of the three directions, and then averaged this value across the three directions. On average, 11.8±1.0 trials were included as low arousal trials, and 15.7±1.7 trials were included as high arousal trials, for each direction at the intermediate spatial frequency (22 FOV, 5 mice). No single direction had fewer than 2 trials at low or high arousal state, and no 3 adjacent directions had fewer than 9 trials in total at low or high arousal state. To further remove noisy boutons for a given spatial frequency, we also required that the DS, AS, BrT and SbC boutons satisfy the criteria for direction-selective, axis-selective, broadly-tuned, or suppressed-by-contrast, respectively. Moreover, for DS, AS, BrT boutons, we required the mean responses across the peak direction and two adjacent directions to be positive at both low and high arousal; for SbC boutons, we required the mean responses to be negative at both low and high arousal. Due to these additional criteria, the total number of boutons used for calculating the modulation index was smaller than the number of boutons used for calculating correlation with pupil area.

For each bouton, we also assessed whether arousal modulation was significant (p<0.05). An N-way ANOVA was used to compare the responses from the aforementioned three directions (direction of maximal response and adjacent directions), at low vs. high arousal.

#### Arousal modulation during the stationary state

The mouse was considered to be stationary during a given trial if, from 1 sec before and 2 sec after onset of visual stimulation, its movement speed didn’t exceed 0.0155 m/s at any moment (i.e. in any frame, with sampling at 15.5 Hz) and the total movement was no more than 0.5 cm, the minimum step size of our rotary encoder. 0.0155 m/s was the minimum unit of running speed during a given frame, and 0.5 cm of movement is equivalent to movement at this minimal speed for 1/3 sec. This criterion ensured that occasionally adjustment of posture during a given trial would still be considered a stationary trial. For analyses of arousal differences during the stationary state, we used the same pupil-based definition of low vs. high arousal trials as above, but we now only included trials that met the additional criterion of reflecting the stationary state. Across the 8 FOV from 4 mice used for assessing arousal modulation during the stationary state, no single stimulus direction had fewer than 3 trials at either the low or high arousal state, and no 3 adjacent directions had fewer than 11 trials in total at either the low or high arousal state. On average, 13.3±1.7 trials were included as low arousal trials, and 12.8±2.5 trials were included as high arousal trials, for each stimulus direction at the intermediate spatial frequency.

##### OnOff preference index

An OnOff preference index was calculated using the averaged response traces to luminance increments (On stimulus) and to luminance decrements (Off stimulus). A positive response to On only, to Off only, or a positive response of equal magnitude to On and Off corresponded to index values of 1, −1 or 0, respectively. Boutons lacking both a significant On response and a significant Off response (see Subsection, ‘Estimation of boutons with significant visual responses’) were not considered.

In order to take into account the dynamics of the evoked On and Off responses, a weighted OnOff preference index was introduced as follows:

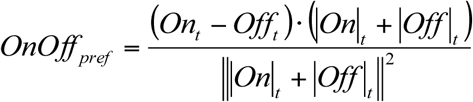

*On_t_* and *Off_t_* were defined as the On and Off response timecourses during the 2-second response window. In this equation, the term (*On_t_* − *Off_t_*) determines the sign of the index at each time point. The dot product of this term with (|*On*|*_t_* + |*Off*|*_t_*) was used to assign a relative weight to each time point according to its summed response magnitude. Then the numerator was normalized to obtain a single preference index between −1 and 1.

A subset of SbC boutons exhibited a positive rebound after transient suppression [13, 14]. The positive values in On and Off response traces from those boutons were set to zero prior to estimating OnOff preference. Moreover, we added a minus sign to the above formula, so that SbC boutons that were significantly suppressed by an On stimulus were defined as On-responsive SbC, while SbC boutons that were significantly suppressed by an Off stimulus were defined as Off-responsive. While this is opposite to our previous definition for SbC cells [2], it increased clarity in this study by enabling consistent interpretation of arousal modulation across bouton classes.

#### Estimation of retinotopic preferences in RGC boutons

##### Retinotopic tuning curve fitting

Two retinotopic tuning curves, which were independently fitted for each bouton, were established, respectively, for tuning along the azimuth and along the elevation axes. Both curves consisted of eight evenly spaced values, each consisting of the average response across trials for a given location in visual space of the oriented bar containing binarized spatiotemporal noise (see above). Tuning curves were approximated using a Gaussian function:

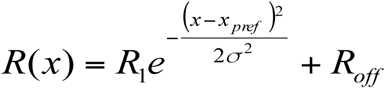

The ΔF/F response, *R*(*x*) varied as a function of the retinotopic stimulus location, *x*. The maximum response, *R*_1_ + *R_off_*, was evoked at *x_pref_*, the preferred retinotopic location. The standard deviation *σ* of the Gaussian was proportional to the receptive field size along this axis. To increase the number of points for fitting from 8 to 15, an interpolation method similar to the one used for direction tuning curves was implemented. As responses were very reliable and well fit, no bootstrapping method was implemented. Fitting was considered significant if 2 out of the 8 directions showed a significant response (see Subsection, ‘Estimation of boutons with significant visual responses’), if the absolute correlation coefficient 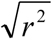 exceeded 0.8, and if the fitted peak amplitude was confirmed to be positive or negative for non-suppressed and for suppressed boutons, respectively.

##### Retinotopic responses of neuropil surrounding RGC axonal boutons

Retinotopic tuning curve fitting was also implemented for the neuropil rings surrounding each bouton, to estimate the local retinotopic preference in the field of view. Each pixel in the field of view was attributed a preferred retinotopic location by first assigning the center of each neuropil ring with a value corresponding to the preferred retinotopic location of that neuropil ring, then dilating by a disk of 10 µm radius from each neuropil center respectively and averaging the preferred retinotopy across overlapping disks. The final pixel-wise estimates of retinotopic preference were obtained by spatial smoothing using an isotropic two-dimensional Gaussian filter with a standard deviation of 3 µm.

### QUANTIFICATION AND STATISTICAL ANALYSIS

Statistical tests were conducted using MATLAB. Non-parametric tests were used for comparing two independent groups (Mann-Whitney-Wilcoxon test), two related groups (Wilcoxon signed-rank test), and multiple independent groups (Kruskal-Wallis test with post-hoc Dunn’s correction), multiple related groups (Friedman test with post-hoc Dunn’s correction). A Kolmogorov-Smirnov test was performed to compare cumulative distributions, p<0.05 was considered significant. Additional details on sample sizes, statistical test, significant levels for each experiment can be found in figure legends, Results and Star Methods. All acquired data were included for analyses.

### DATA AND SOFTWARE AVAILABILITY

Requests for analyses and raw data on calcium imaging results may be made to the Lead Contact, Mark L. Andermann.

